# An ERAD-independent role for rhomboid pseudoprotease Dfm1 in mediating sphingolipid homeostasis

**DOI:** 10.1101/2022.07.30.502165

**Authors:** Satarupa Bhaduri, Analine Aguayo, Yusuke Ohno, Marco Proietto, Jasmine Jung, Isabel Wang, Rachel Kandel, Narinderbir Singh, Ikran Ibrahim, Amit Fulzele, Eric Bennett, Akio Kihara, Sonya E. Neal

**Affiliations:** School of Biological Sciences, Department of Cell and Developmental Biology, University of California San Diego, La Jolla, CA 92093; Laboratory of Biochemistry, Faculty of Pharmaceutical Sciences, Hokkaido University, Kita 12-jo, Nishi 6-chome, Kita-ku, Sapporo 060-0812, Japan

## Abstract

Nearly one-third of nascent proteins are initially targeted to the endoplasmic reticulum (ER) where they are correctly folded and assembled before being delivered to their final cellular destinations. To prevent the accumulation of misfolded membrane proteins, <u>ER</u>-<u>a</u>ssociated-<u>d</u>egradation (ERAD) removes these clients from the ER membrane to the cytosol in a process known as retrotranslocation. Our recent work demonstrates that rhomboid pseudoprotease, Dfm1, is involved in the retrotranslocation of ubiquitinated integral membrane ERAD substrates. To survey for potential interaction partners of Dfm1, we performed protein-proximity labeling by BioID (proximity-dependent <u>bio</u>tin <u>id</u>entification) followed by mass spectrometry and identified several interacting proteins known to play a role in the sphingolipid biosynthesis pathway. Specifically, we found that Dfm1 physically interacts with the SPOTS complex, which is composed of serine palmitoyltransferase (SPT) enzymes and accessory components and is critical for catalyzing the first rate-limiting step of the sphingolipid biosynthesis pathway. We demonstrate for the first time that Dfm1 has a role in ER export, a function that is independent of Dfm1’s canonical ERAD retrotranslocation function. Specifically, we show that loss of Dfm1 results in the accumulation of phosphorylated Orm2 at the ER, suggesting a novel role for Dfm1 in controlling Orm2 export from the ER and its subsequent degradation by EGAD. Moreover, recruitment of Cdc48 by Dfm1, which is critical for its role in ERAD retrotranslocation, is dispensable for Dfm1’s role in ER export. Given that the accumulation of human Orm2 homologs, ORMDLs, are associated with many maladies, our study serves as a molecular foothold for understanding how dysregulation of sphingolipid metabolism leads to various diseases.

## INTRODUCTION

The endoplasmic reticulum (ER) carries out a vast range of functions including protein synthesis and transport, protein folding, lipid and steroid synthesis, carbohydrate metabolism, and calcium storage. Almost all eukaryotic membrane and secreted proteins are co-translationally imported into the ER where they are subsequently folded (Sicari et al., 2019; Wang and Dehesh, 2018). Proteins frequently fail to fold or assemble properly, at which point they are eliminated by ER-Associated-Degradation (ERAD) (Mehrtash and Hochstrasser, 2019; Ruggiano et al., 2014; Sun and Brodsky, 2019).

ERAD describes a range of pathways that target and ubiquitinate a large repertoire of secretory and membrane substrates for proteasomal degradation. ERAD substrates are classified according to the location of their lesions and are referred as ERAD-L (lesion in luminal domain), ERAD-M (lesion within the transmembrane domain), and ERAD-C (lesion in the cytosolic domain). The HMG-CoA reductase degradation (HRD) pathway utilizes the E3 ligase, Hrd1, to target ERAD-M and ERAD-L substrates, and the degradation of alpha 2 (DOA) pathway utilizes the E3 ligase, Doa10, to target ERAD-C substrates (Carvalho et al., 2006; Foresti et al., 2013; Hampton et al., 1996; Hiller et al., 1996; Laney and Hochstrasser, 2003). Moreover, unassembled ER subunits escaping to the inner nuclear membrane (INM) are targeted by the Asi complex (Foresti et al., 2014; Natarajan et al., 2020). A common theme for all ERAD pathways is the removal or retrotranslocation of ubiquitinated substrates from the ER membrane or INM followed by degradation by the proteasome (Hampton and Sommer, 2012). Retrotranslocation has been well-characterized in *S. cerevisiae*, with two derlins, Der1 and Dfm1, serving as major mediators of retrotranslocation for ERAD-L and ERAD-M substrates, respectively (Neal et al., 2018; Wu et al., 2020). Moreover, previous structural studies suggest that the multi-membrane spanning yeast E3 ligases, Hrd1 and Doa10, function as channels for the retrotranslocation of luminal and single-spanning membrane substrates, respectively (Schmidt et al., 2020; Wu et al., 2020). No analogous channel for multi-spanning membrane substrates had been determined, until Neal and colleagues identified the yeast derlin, Dfm1, as a major retrotranslocation factor for a subset of membrane substrates (Neal et al., 2018).

Dfm1 is an ER-resident multi-spanning membrane protein and is classified as a rhomboid pseudoprotease. Recently, we showed that Dfm1 utilizes its conserved rhomboid protein residues for substrate engagement and its lipid thinning properties to allow retrotranslocation of multi-spanning membrane substrates (Nejatfard et al., 2021). To identify interacting partners of Dfm1 that may assist with retrotranslocation, we employed proximity-based labeling followed by mass spectrometry. Remarkably, we identified several proteins enriched with Dfm1, which are known to play a role in the sphingolipid biosynthesis pathway. Sphingolipids constitute a major class of lipids defined by their amino-alcohol backbone with mainly eighteen-carbon and are synthesized in the ER from non-sphingolipid precursors (Hannun and Obeid, 2018). Modification of this basic structure gives rise to the vast family of sphingolipids, which have essential roles in cell signaling and function. Serine palmitoyltransferase (SPT) is the first rate-limiting enzyme in the de novo synthesis of sphingolipids, and its sole function is to catalyze the initial step in sphingolipid biosynthesis by converting serine and palmitoyl-CoA into a sphingolipid precursor, 3-keto-sphinganine (Hanada, 2003). SPT is essential for the viability of all eukaryotic cells, and mutations of SPT are linked to hereditary sensory neuropathy type 1 (HSAN1) and early onset amyotrophic lateral sclerosis (ALS) (Bode et al., 2015; Mohassel et al.). Accordingly, SPT serves as the key point for regulation of sphingolipid biosynthesis. SPT forms the SPOTS complex, comprised of Orm1 and Orm2 (members of the orosomucoid (ORM) gene family), Tsc3, and Sac1.The SPOTS complex is highly conserved from yeast to mammals (Breslow et al., 2010). Functionally, Orm1, Orm2, and phosphoinositide phosphatase Sac1 are evolutionarily conserved negative regulators of SPT, while Tsc3 is a positive regulator.

Previous studies have demonstrated that levels of several SPOT complex members and sphingolipid biosynthesis enzymes are regulated through protein degradation pathways in order to control sphingolipid levels. For example, the TORC2-Ypk1 signaling axis phosphorylates Orm2, triggering its export from the ER to the Golgi, where it is selectively ubiquitinated by the Dsc complex before being retrotranslocated and degraded by the cytosolic proteasome. This pathway is known as ER Golgi-Associated Degradation (EGAD)(Schmidt et al., 2019, 2020). Another study analyzing systematic turnover of proteins in yeast revealed that several enzymes and regulators involved in the de novo sphingolipid biosynthesis pathway are degraded in separate organelles, such as the Golgi and vacuole (Christiano et al., 2020). Although many enzymes and regulators associated with sphingolipid biosynthesis reside in the ER, an ER-localized regulator for sphingolipid homeostasis has not yet been identified. In this study, we report a novel role for the ER-resident Dfm1 in maintaining sphingolipid homeostasis. We find that Dfm1 physically and genetically interacts with SPOTS complex components. This includes a genetic interaction with TSC3, a positive regulator of SPT, whose function is essential for stimulating SPT activity at 37°C. Specifically, loss of Dfm1 rescues the growth lethality of *tsc3*Δ cells by increasing ceramide and complex sphingolipid levels. DFM1 also genetically interacts with ORM1, a negative regulator of SPT activity, in which *orm1*Δ*dfm1*Δ cells have an exacerbated growth defect due to increased flux in sphingolipid biosynthesis. Finally, we provide the first evidence that Dfm1 is required for Orm2 degradation, a function that is independent of Dfm1’s classical ERAD-M retrotranslocation function. We confirm the independence of Dfm1’s ERAD function and demonstrate that the EGAD-client, Orm2, does not require ERAD nor Inner Nuclear Membrane Associated Degradation (INMAD) pathways, which is in agreement with an earlier study (Schmidt et al., 2019). To better understand the role of Dfm1 in Orm2 degradation, we show that loss of Dfm1 results in accumulation of phosphorylated Orm2 at the ER, suggesting a novel role for Dfm1 in controlling Orm2 export from the ER and its subsequent degradation by EGAD. Although Dfm1 mediates Orm2 export from the ER, it does not directly function with COPII dynamics and trafficking. Overall, our work identifies the highly conserved derlin Dfm1 as a critical mediator of sphingolipid homeostasis and provides a new therapeutic target for maladies associated with dysregulation in sphingolipid homeostasis.

Overall, this study demonstrates a novel role for the highly conserved derlins in regulating sphingolipid metabolism. Given that the accumulation of human Orm1/2 paralogs, ORMDLs, are associated with many maladies, our study serves as a molecular foothold for understanding how dysregulation of sphingolipid metabolism leads to various diseases.

## RESULTS

### Derlin Dfm1 interacts with members of the sphingolipid biosynthetic pathway

To identify potential Dfm-1 interacting proteins, Proximity-dependent biotin identification (BioID) was employed. Briefly, BirA-3xFLAG was fused to Dfm1 to survey for potential interacting partners (Fig. 1A). Because Dfm1 included an added BirA-3xFLAG epitope at the C-terminus, we wished to confirm that the tag did not affect the expression and function of Dfm1. To this end, tagged-DFM1 was placed under a galactose inducible promoter (GAL_pr_) and cells expressing GAL-driven Dfm1-BirA-3xFLAG were grown in the presence of 2% galactose. Under these conditions, induced expression of Dfm1 was observed at the expected molecular weight of ∼60 kDa (Fig. 1B). BirA-3xFLAG alone expressed at both the expected size of ∼25 kDa and at a larger size, which most likely represents BirA aggregates (marked by asterisks, Fig. 1B). To test whether Dfm1-BirA-3xFLAG function is still intact, we performed cycloheximide (CHX)-chase of a well-characterized Dfm1 substrate, Hmg2(Hampton et al., 1996). We observed Hmg2-GFP degradation upon addback of Dfm1-BirA-3xFLAG while Hmg2-GFP degradation was stabilized with both empty vector and BirA-3xFLAG alone (Fig. 1C). To validate the identification of Dfm1 interactors via biotinylation, cells were treated with biotin and biotinylated proteins were enriched with streptavidin beads. As expected, the ATPase Cdc48, which has previously been shown to bind directly to Dfm1 (Neal et al., 2018; Sato and Hampton, 2006), was enriched in the biotin-treated samples, whereas neither Cdc48 nor Dfm1 were enriched in untreated or BirA-3xFLAG alone cells (Fig. 1D). Next, proteins that were enriched with streptavidin beads were digested to obtain tryptic peptides and analyzed by LC/MS/MS. Quantified proteins were mapped on volcano plots based on the significance and the ratio between Dfm1-BirA-3xFLAG and controls. High-confidence interacting proteins were identified using a modified version of CompPASS (Fig. 1E) (Zuzow et al., 2018). By applying gene ontology (GO) enrichment analyses for the sets of Dfm1 interacting proteins identified, we found GO terms related to “Ceramide Metabolic Process” to be the most enriched (Fig. 1F). The interactions were validated by the presence of several ERAD components (Hrd1, Cdc48, proteasome subunits: Rpt2 and Pre9). Interestingly, closer analysis revealed unexpected interactions with SPOTS complex members (Orm1, Tsc3, and Lcb2). Taken together, these results suggest that our data have a high level of confidence and represent a rich source of Dfm1 interactome proteins, which include members of the sphingolipid biosynthesis pathway.

**Fig. 1.**
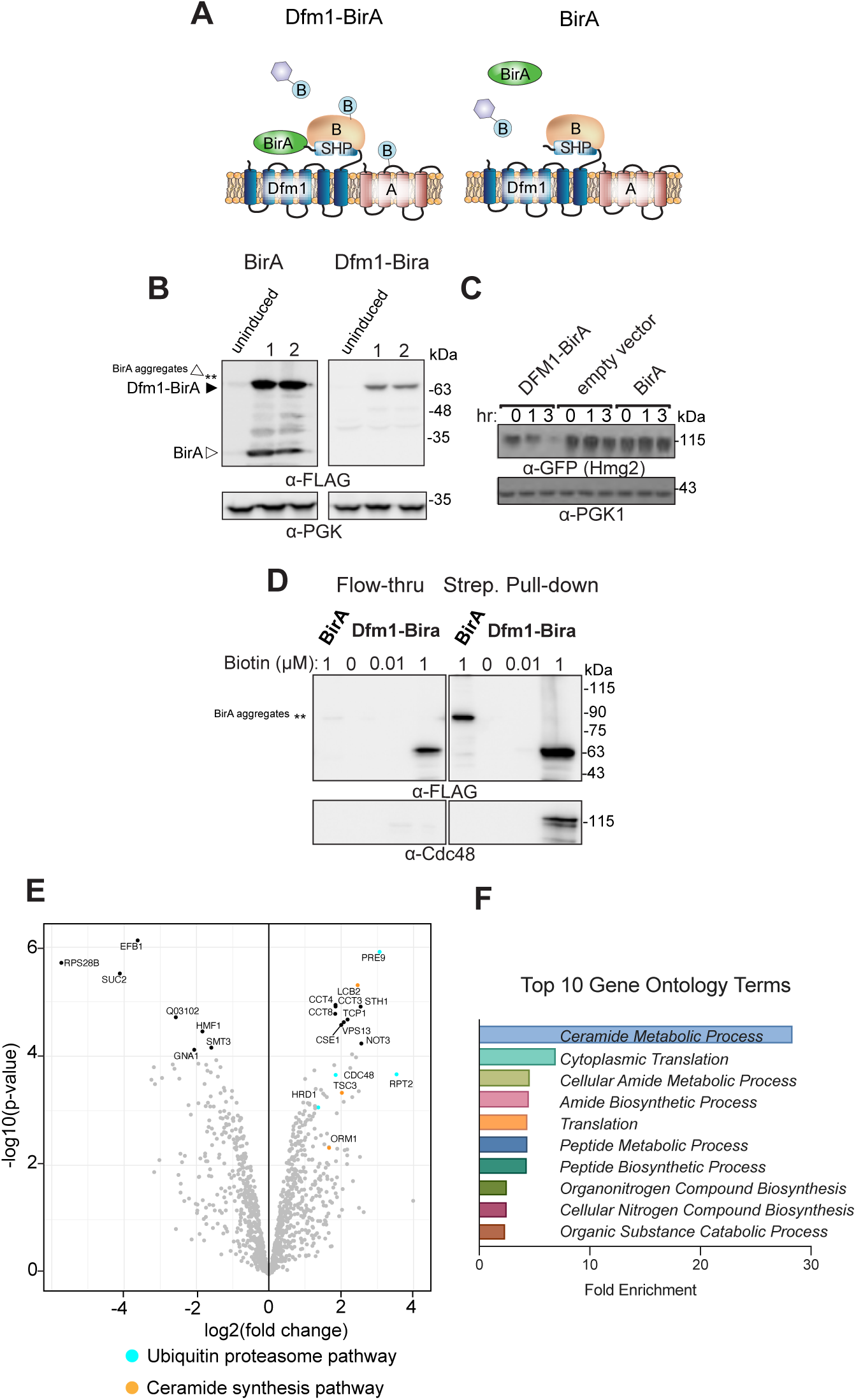
BioID proximity-based labeling to identify interaction partners of Dfm1. **(A)** Schematic of Dfm1 fused with a biotin ligase, BirA, at the C-terminus along with un-tagged Dfm1.Cartoon representation of the labeling of rhomboid pseudoprotease Dfm1 with the biotin ligase, BirA. (**B**) GALpr-Dfm1-BirA-Flag and GALpr-BirA-Flag levels were measured by western blotting with α-FLAG at 0 (uninduced) vs. 5 hours post-galactose induction. **(C)** Dfm1-BirA is still functional and able to degrade Hmg2-GFP. *dfm1*Δ+Hmg2-GFP strains containing DFM1-BIRA, empty vector, or BIRA only addbacks were grown to log phase and degradation was measured by CHX. After CHX addition, cells were lysed at the indicated times and analyzed by SDS-PAGE and immunoblotted for Hmg2-GFP with α-GFP. (**D)** Yeast strains expressing Dfm1-Bira and BirA only negative control were incubated with different amounts of biotin: 0, 0.1, and1 mM. *dfm1*Δ+Hmg2-GFP. Microsomes were isolated from triplicates of each strain and subjected to streptavidin pulldown. Flow-through and pull-down fractions were detected by western blotting for Dfm1-BirA with α-Flag and Cdc48 with α-Cdc48 antibodies. **(D)** A volcano plot showing enrichment versus significance of proteins identified in Dfm1-BirA experiments relative to control (BirA only) experiments. Components with enrichment of at least 1.5 fold and p>.05 are ERAD components in blue (Hrd1, Cdc48, Pre9, and Rpt2) and sphingolipid biosynthesis in orange (Lcb2, Tsc3, and Orm1). **(E)** Top 10 gene ontology (GO) terms and their enrichment factor for the set of genes with the highest significance and fold enrichment.

To validate the interaction of Dfm1 with SPOTS complex members, we performed co-immunoprecipitation (co-IP). Cells co-expressing Dfm1-GFP and members of the SPOTS complex (Lcb1-RFP, and Orm2-RFP) were subjected to immunoprecipitation via GFP Trap. Notably, Lcb1-RFP and Orm2-RFP co-immunoprecipitated with Dfm1-GFP, whereas no detectable association was seen in control cells without Dfm1-GFP (Fig. 2A). These interactions were also validated by fluorescence microscopy, in which the majority of Dfm1-GFP co-localized with Lcb1-RFP and Orm2-RFP at the ER (Fig. 2B). Hence, Dfm1 binds directly to SPOTS complex members at the ER.

**Fig. 2.**
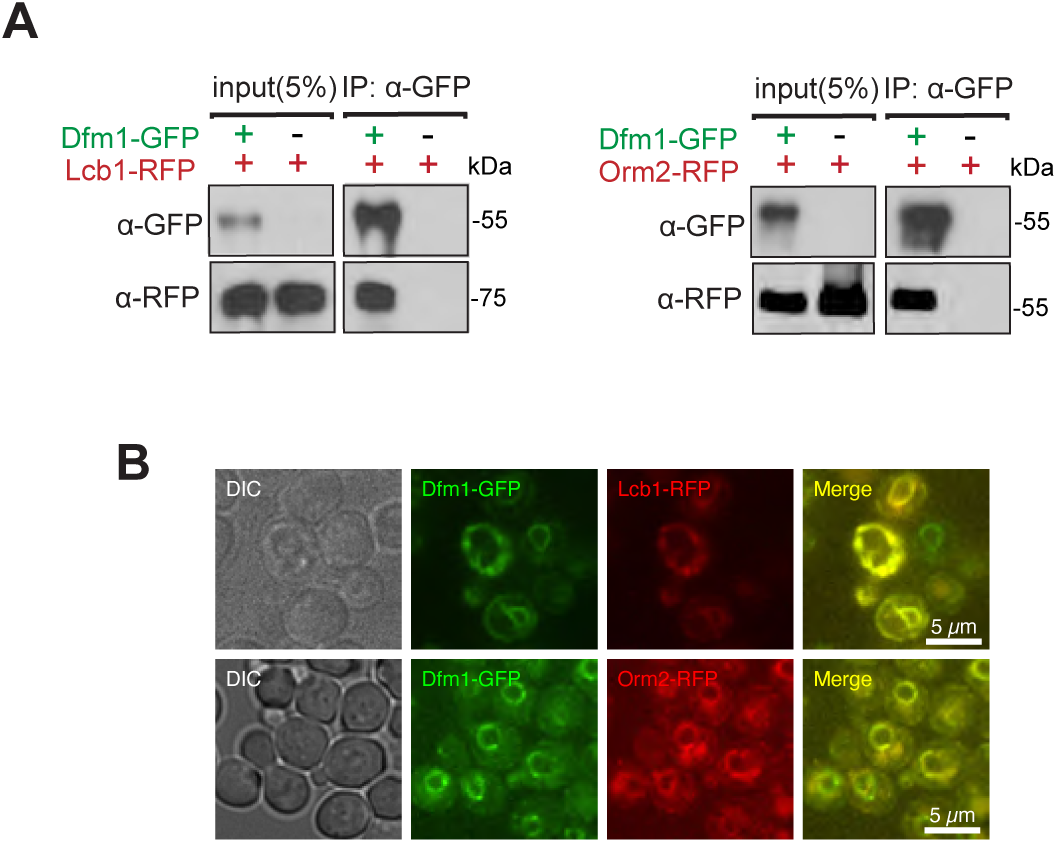
Dfm1 colocalizes and binds to SPOTS complex proteins. **(A)** Dfm1-GFP binding to Lcb2-RFP and Orm2-RFP were analyzed by co-IP. As negative control, cells not expressing Dfm1-GFP were used. (B) Dfm1-GFP colocalizes with Lcb1-RFP and Orm22-RFP. Strains were grown to mid-exponential phase in minimal media GFP and RFP fluorescence was examined on an AxioImager.M2 fluorescence microscope using a 100x objective and 28HE-GFP or 20HE-rhodamine filter sets (Zeiss).

### DFM1 genetically interacts with TSC3

We next examined whether Dfm1 genetically interacts with SPOTS complex members. The SPOTS complex consists of the SPT enzymes, Lcb1 and Lcb2, and the smaller subunit, Tsc3, which has been required to positively regulate SPT at high temperatures (Gable et al., 2000). Furthermore, SPT activity is negatively regulated by two yeast paralogs, Orm1 and Orm2, through direct interactions, and by Sac1, which negatively regulates SPT through an unknown mechanism (Breslow et al., 2010; Han et al.). To survey for gene interactions, we generated double mutant yeast strains of *dfm1*Δ along with respective SPOTS complex members and performed serial dilution growth assays to test whether double knockout cells confer any distinct growth phenotypes compared with WT and single knockouts. To test the involvement of essential enzymes Lcb1/Lcb2 and non-essential regulator Sac1, we utilized *Lcb1-DaMP*, *Lcb2-DamP*, and *sac1*Δ mutants and observed no genetic interactions, since growth of *dfm1*Δ*Lcb1-DaMP*, *dfm1*Δ*Lcb2-DaMP*, and *dfm1*Δ*sac1*Δ was similar to that of WT cells at 25°C, 30°C, and 37°C (Fig. S1A). A small subunit of the SPT, Tsc3, directly interacts with Lcb1/Lcb2 to stimulate their activity and increase synthesis of the sphingolipid precursor, 3-ketosphinganine. The stimulatory function of Tsc3 is essential at the higher temperature where the *tsc3*Δ temperature-sensitive phenotype is lethal due to lack of phytosphingosine (PHS) production ((Gable et al., 2000) and Fig. 3A). In line with this observation, we observed a growth defect and lethality from *tsc3*Δ cells at 30°C and 37°C, respectively (Fig. 3A, *filled triangle*), and rescue of lethality when PHS was supplied to *tsc3*Δ cells (Fig. 3B, *right panel; filled triangle*). Remarkably, removal of DFM1 in this background – *dfm1*Δ*tsc3*Δ – completely rescued the lethality at 37°C (Fig. 3A, *open circle*). Thus, removal of DFM1 suppresses *tsc3*Δ lethality at 37°C.

**Fig. 3.**
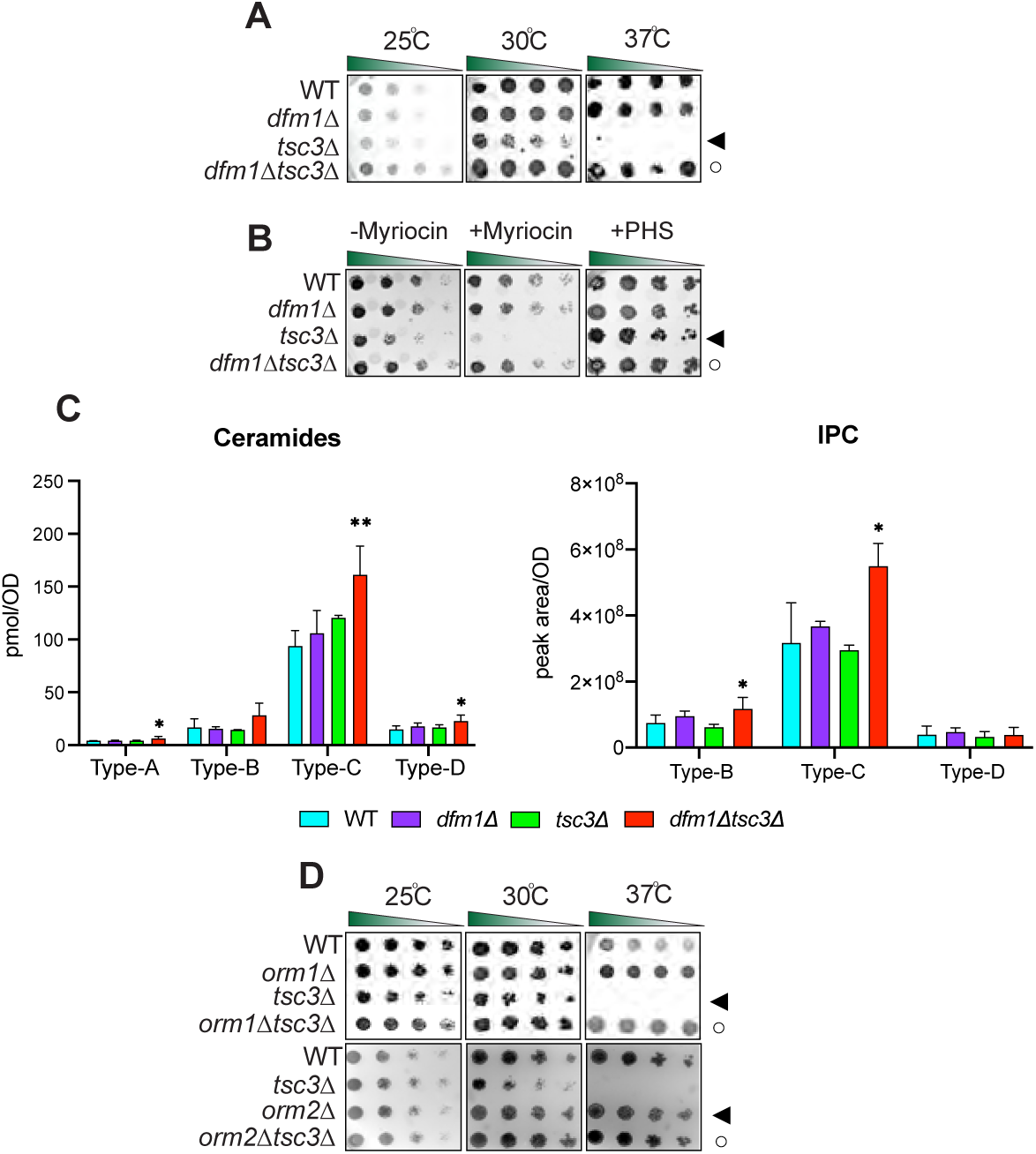
Dfm1 genetically interacts with Tsc3. **(A)** Indicated strains were spotted 5-fold dilutions on synthetic complete (SC) plates in triplicates, and plates were incubated at room temperature, 30°C, and 37°C (n=3). WT, *dfm1*Δ, *tsc3*Δ, and *dfm1*Δ*tsc3*Δ were compared for growth in the dilution assay. **(B)** *dfm1*Δ*tsc3*Δ confers resistance to myrocin. WT, *dfm1*Δ, *tsc3*Δ, and *dfm1*Δ*tsc3*Δ strains were grown to log-phase in YPD medium, and 5-fold serial dilutions of cultures were spotted on (SC) plates containing either drug vehicle alone, 1 µM of myriocin and 10 μM of PHS. **(C)** WT, *dfm1*Δ, *tsc3*Δ, and *dfm1*Δ*tsc3*Δ cells were grown to log-phase at 30°C and lipids were extracted and subjected to LC-MS/MS. A- B- C- & D-Type ceramides containing C16, C18, C20, C22, C24, and C26 fatty acid (left graph) and B- C- & D-type IPCs containing C24 and C26 fatty acid (right graph) were measured. Values represent the means±s.d.s of three independent experiments. Statistically significant differences compared to WT cells are indicated (Dunnett’s test; *P<0.05, \*\**P*<0.01). **(D)** Indicated strains were spotted 5-fold dilutions on SC plates in triplicates, and plates were incubated at room temperature, 30°C, and 37°C (n=3). *Upper panel:* WT, *orm1*Δ, *tsc3*Δ, *and orm1*Δ*tsc3*Δ were compared for growth by dilution assay. *Middle panel:* WT, *orm2*Δ, *tsc3*Δ, *and orm2*Δ*tsc3*Δ were compared for growth by dilution assay. *Bottom panel:* WT, *orm1*Δ, *orm2*Δ, *tsc3*Δ, *and orm1*Δ*orm2*Δ*tsc3*Δ, were compared for growth by dilution assay.

### dfm1Δtsc3Δ cells have increased steady-state levels of ceramides and complex sphingolipids

We predicted that removal of DFM1 was able to reverse the temperature-sensitive lethality in *tsc3*Δ as a result of increased production of sphingolipid precursors. Notably, myriocin is a potent inhibitor of SPT, the first committed step in the sphingolipid biosynthesis pathway, and treatment with myriocin reduces sphingolipid levels in both *S. cerevisiae* and mammals (Breslow, 2013). Because SPT activity is essential, myriocin treatment exacerbates growth due to decreased flux in the sphingolipid biosynthesis pathway. We therefore wanted to test whether *dfm1*Δ*tsc3*Δ cells are resistant to myriocin inhibition. To this end, a sublethal dose of myriocin was used in the serial growth assay to reduce sphingolipid synthesis without impairing cell growth. As expected, *tsc3*Δ cells were sensitive to myriocin treatment at 30°C since these cells already have decreased sphingolipid levels (Fig. 3B, *left panel; filled triangle*). By contrast, the *dfm1*Δ*tsc3*Δ cells were resistant to myriocin treatment, suggesting that these cells have higher levels of sphingolipids (Fig. 3B, *left panel; open circle*). Indeed, when grown at 30°C, lipidomic analysis demonstrated that *dfm1*Δ*tsc3*Δ cells significantly produced higher levels of ceramides (Type A, C, & D) and complex sphingolipid inositolphosphorylceramide (IPC) (Type B & C) in comparison to WT cells (Fig. 3C). Altogether, *dfm1*Δ*tsc3* cells appear to produce higher steady-state levels of ceramide and complex sphingolipids.

### DFM1 genetically interacts with ORM1

Because removal of DFM1 leads to higher levels of ceramides and complex sphingolipids in *tsc3*Δ cells, we hypothesized that Dfm1 is antagonizing the sphingolipid biosynthesis pathway. If this hypothesis is correct, *dfm1*Δ cells should phenocopy both *orm1*Δ and *orm2*Δ cells, which are established negative regulators of the SPT enzymes, in the growth assays. To test this hypothesis, *orm1*Δ*tsc3*Δ and *orm2*Δ*tsc3*Δ cells were generated and employed in the growth assays at 25°C, 30°C, and 37°C (Fig. 3D). Under these conditions, both *orm1*Δ and *orm2*Δ phenocopied *dfm1*Δ; both *orm1*Δ*tsc3*Δ and *orm2*Δ*tsc3*Δ cells were able to rescue the temperature-sensitive lethality displayed by *tsc3*Δ cells (Fig. 3D). Because *dfm1*Δ cells phenocopy both *orm1*Δ and *orm2*Δ cells, we next examined whether DFM1 genetically interacts with either ORM1 or ORM2. Although no growth defect was observed for *dfm1*Δ*orm2*Δ cells, we did observe a growth defect in *dfm1*Δ*orm1*Δ cells at room temperature, 30°C, and 37°C, suggesting that DFM1 functions with ORM1 in a parallel pathway (Fig. 4A, *filled triangle*). Furthermore, lipidomic analysis confirmed that *dfm1*Δ*orm2*Δ cells showed no increase in ceramides compared with WT cells. This in contrast to *dfm1*Δ*orm1*Δ cells where there were significant changes in ceramides levels compared to WT cells (discussed below), further supporting the hypothesis that *dfm1*Δ genetically interacts with *orm1*Δ but not *orm2*Δ (Fig. 4B).

**Fig. 4.**
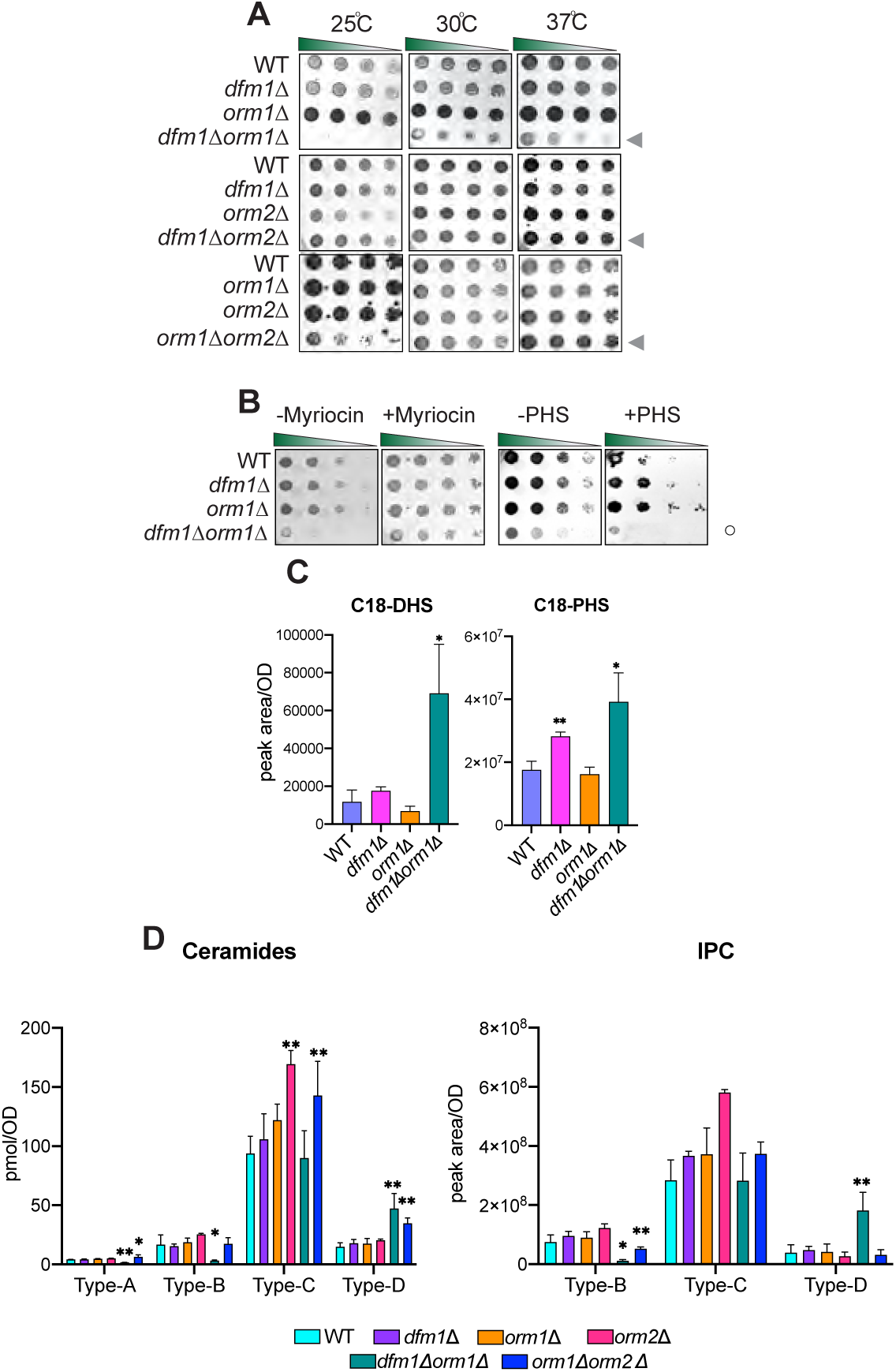
Dfm1 genetically interacts with Orm1. **(A)** Indicated strains were spotted 5-fold dilutions on SC plates in triplicates, and plates were incubated at room temperature, 30°C, and 37°C (n=3). *Upper panel:* WT, *dfm1*Δ, *orm1*Δ, *and dfm1*Δ*orm1*Δ were compared for growth by dilution assay. *Middle panel:* WT, *dfm1*Δ, *orm2*Δ, *and orm2*Δ*tsc3*Δ were compared for growth by dilution assay. *Bottom panel:* WT, *dfm1*Δ, *orm1*Δ, *orm2*Δ, *and orm2*Δ, and *orm1*Δ*orm2*Δ were compared for growth by dilution assay. **(B)** *dfm1*Δ*orm1*Δ confers resistance to myriocin and sensitivity to PHS. WT, *dfm1*Δ, *orm1*Δ, and *dfm1*Δ*orm1*Δ strains were grown to log-phase in SC medium, and 5-fold serial dilutions of cultures were spotted on YPD plates containing either drug vehicle alone, 1 mM of myriocin and 10 μM of PHS. Plates were incubated at room temperature and photographed after 3 days. **(C)** WT, *dfm1*Δ, *orm1*Δ, and *orm1*Δ*dfm1*Δ cells were grown to log-phase at 30°C and lipids were extracted and subjected to LC-MS/MS. C18 PHS and DHS levels were measured as described in Methods. Values represent the means±s.d.s of three independent experiments. Statistically significant differences compared to WT cells are indicated (Dunnett’s test; *P<0.05, \*\**P*<0.01). **(D)** WT, *dfm1*Δ, *orm1*Δ, *orm2*Δ, *dfm1*Δ*orm1*Δ, and *orm1*Δ*orm2*Δ cells were grown to log-phase at 30°C and lipids were extracted and subjected to LC-MS/MS. A- B- C- & D-Type ceramides containing C16, C18, C20, C22, C24, and C26 fatty acid (left graph) and B- C- & D-type IPCs containing C24 and C26 fatty acid (right graph) were measured. Results are the average ± SD of ceramide or IPC levels from three independent colonies (*** *p* < 0.001, *n* = 3; N.·S, not significant, *n* = 3). Results are representative of multiple independent experiments.

Previous studies have demonstrated that cells lacking ORM1 and ORM2 exhibit a growth defect (Fig. 4A, *bottom panel*) due to an increased flux in de novo sphingolipid synthesis and the knockout cells were more resistant to myriocin inhibition (Breslow et al., 2010; Han et al.). Given the growth defect seen in *dfm1*Δ*orm1*Δ cells, we reasoned that the flux in sphingolipid synthesis should be similarly increased. Notably, *dfm1*Δ*orm1*Δ cells were resistant to myriocin treatment (Fig. 4B, *left panel; open circle*) and sensitive to exogenously-added PHS (Fig. 4B, *right panel; open circle*), since *dfm1*Δ*orm1*Δ cells already exhibit higher levels of sphingolipids. Both dihydrosphingosine (DHS) and phytosphingosine (PHS) are early precursors of the sphingolipid biosynthesis pathway and are derivatives of long-chain bases (LCBs). Lipidomic analysis via mass spectrometry showed that C18-DHS levels were significantly higher in *dfm1*Δ*orm1*Δ cells than in WT cells. Also, C18-PHS levels were significantly higher in both *dfm1*Δ and *dfm1*Δ*orm1*Δ cells in comparison to WT cells, suggesting there is increased flux in sphingolipid biosynthesis in *dfm1*Δ*orm1*Δ cells (Fig. 4C). In contrast, the levels of ceramides and complex sphingolipids varied in *dfm1*Δ*orm1*Δ cells. There were higher levels of ceramide and complex sphingolipids (Type D) and lower levels of ceramide (Type A,B,&C) and complex sphingolipids (Type B) in *dfm1*Δ*orm1*Δ cells in comparison to WT cells (Fig. 4E). Notably, *orm1*Δ*orm2*Δ control cells also exhibited similar fluctuating levels of the varying types of ceramides and complex sphingolipids (Fig. 4E). Despite varying levels of ceramides and complex sphingolipids, *dfm1*Δ*orm1*Δ cells have higher LCB levels and are resistant to myriocin treatment, which suggests that the major physiological effect of *dfm1*Δ*orm1*Δ cells is from increased SPT activity (Fig. 4B&C).

### Orm2 is targeted by Dfm1 for degradation

Given the myriad biological processes carried out by sphingolipids, it is not surprising that disruptions to sphingolipid homeostasis have deleterious effects and must be tightly regulated. One possible mode of regulation is through regulated degradation of key enzymes and regulators of sphingolipid biosynthesis in a manner analogous to the regulated degradation of Orm2 by EGAD to establish sphingolipid homeostasis. We therefore tested whether key enzymes or regulators within the sphingolipid biosynthesis pathway are targeted for Dfm1-mediated degradation by performing cycloheximide (CHX)-chase assays on candidate substrates (Lcb1, Lcb2, Orm1, Orm2, Sac1, Tsc3, Ypk1 and Tsc10), which function in either SPT synthesis or regulation (Fig. S2A). Of these, Orm2 was rapidly degraded in wild-type strains and its degradation was completely prevented in *dfm1*Δ cells (Fig. 5A). The yeast paralog of Dfm1, Der1, has a strong broad role in retrotranslocating ERAD-L substrates (Wu et al., 2020). We therefore directly tested the role of Der1 in Orm2 degradation using the CHX-chase assay and found that in both WT and *der1*Δ cells, Orm2 was still degraded (Fig. 5B). These results imply that the degradation of Orm2 is specifically dependent on derlin Dfm1 and not Der1.

**Fig. 5.**
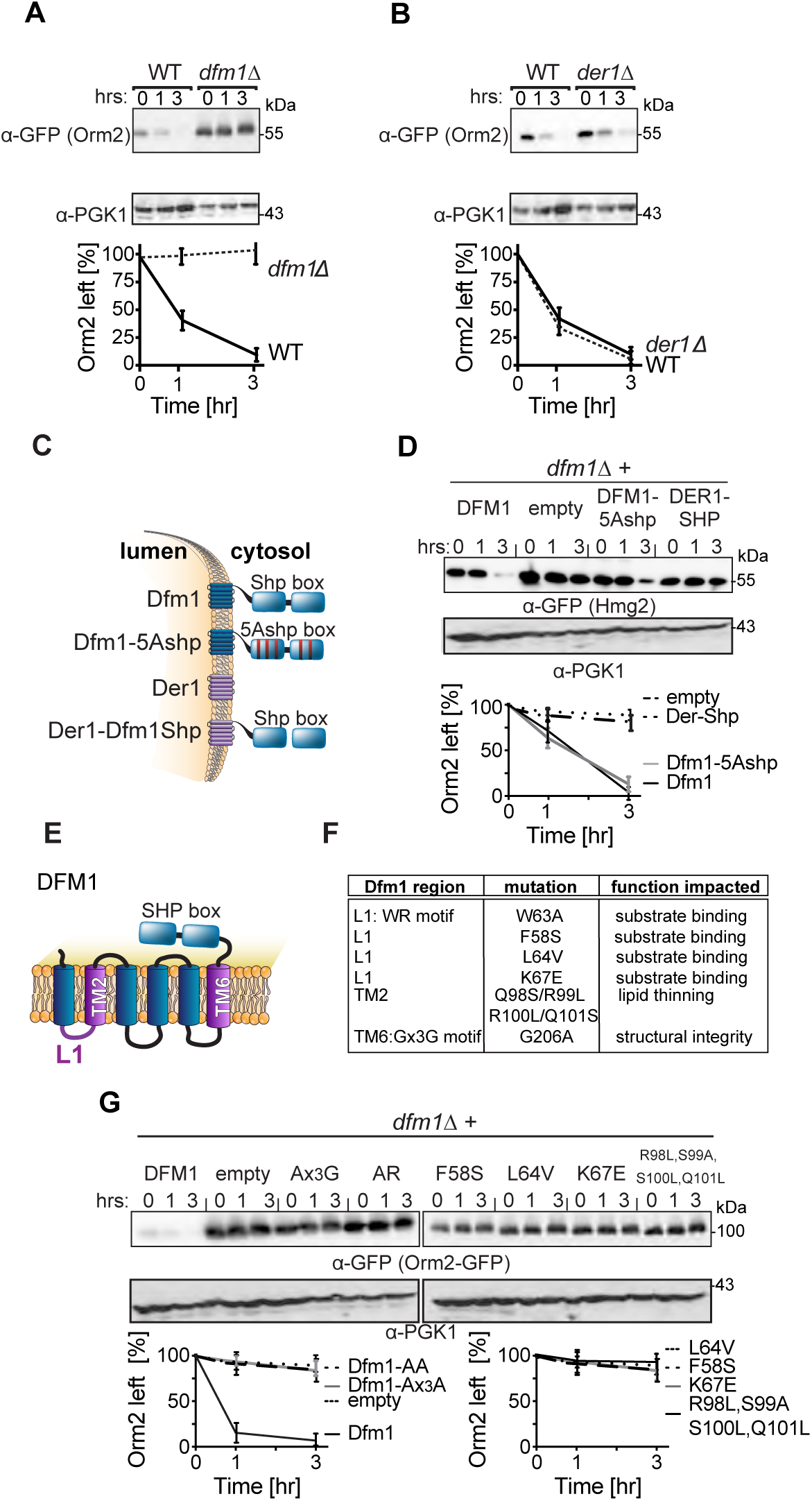
Dfm1 targets Orm2 for degradation. **(A)** Degradation of Orm2 depends on Dfm1 and not Der1. The indicated strains expressing Orm2-RFP were grown into log phase and degradation was measured by cycloheximide chase (CHX). After CHX addition, cells were lysed at the indicated times, and analyzed by SDS-PAGE and immunoblotted for Orm2-RFP with α-RFP. Band intensities were normalized to PGK1 loading control and quantified by ImageJ. t=0 was taken as 100% and data is represented as mean ± SEM from at least three experiments. **(B)** Same as (A) except degradation of Orm2-RFP was measured in WT and *der1*Δ cells. **(C)** Depiction of Dfm1, Der1, Dfm1-5Ashp, and Der1-Shp. Dfm1 and Der1 are ER-localized membrane proteins with six transmembrane domains (Greenblatt et al., 2011). Unlike Der1, Dfm1 has an extended cytoplasmic tail containing two SHP boxes. **(D)** Dfm1’s SHP box is not required for degradation of Orm2-RFP. In the indicated strains, degradation of Orm2-RFP was measured by CHX-chase assay. Cells were analyzed by SDS-PAGE and immunoblotted for Orm2-RFP with α-RFP. **(E)** Depiction of Dfm1, which highlights L1, TM2, TM6, and its SHP box domain. **(F)** Table indicating the location and specific function that is impaired for retrotranslocation-deficient Dfm1 mutants (Nejatfard et al., 2021). **(G)** Dfm1’s WR motif, GxxxG motif, substrate binding and lipid thinning function are required for degradation of Orm2-RFP. In the indicated strains, degradation of Orm2-RFP was measured by CHX-chase assay. Cells were analyzed by SDS-PAGE and immunoblotted for Orm2-RFP with α-RFP.

### Derlin Dfm1’s Cdc48 recruitment function is not required for Orm2 degradation

We have previously identified specific motifs and residues of Dfm1 that are critical for its ERAD retrotranslocation function. Accordingly, we wished to test the importance of these motifs/residues for Orm2 degradation by performing CHX-chase assays. Dfm1 possesses a unique C-terminal SHP box motif, which recruits the ATPase, Cdc48, directly to the ER surface (Neal et al., 2018). Cdc48 functions as an energy source for membrane substrate retrotranslocation and as a retrochaperone where it acts to maintain the solubility of retrotranslocated membrane substrates prior to proteasome degradation (Neal et al., 2017). We previously demonstrated that mutations within the SHP box, Dfm1-5Ashp, ablates Cdc48 recruitment and the retrotranslocation function of Dfm1 (Fig. 5C) (Neal et al., 2018). We also demonstrated that a Der1-SHP chimera, which consists of Der1, the paralog of Dfm1, fused to the cytoplasmic SHP tail of Dfm1, supports Cdc48 recruitment via binding of Cdc48 to the chimera’s SHP tail, but does not support retrotranslocation through Der1’s transmembrane segment (Fig. 5C) (Neal et al., 2018). We utilized these retrotranslocation-deficient variants in our CHX-chase assay to test whether Dfm1’s Cdc48 recruitment function is required for Orm2 degradation. Addback of Der1-SHP in *dfm1*Δ cells impaired Orm2 degradation whereas Dfm1-5Ashp addback still enabled Orm2 degradation (Fig. 5D). These results suggest that recruitment of Cdc48 by Dfm1 is dispensable for Orm2 degradation. In addition, the inability of Der1-SHP to facilitate degradation of Orm2 implies the involvement of additional residues within the transmembrane segments of Dfm1.

Dfm1 contains the highly conserved WR motif in loop 1 (L1) and a Gx3G motif in transmembrane 6 (TM6). Both motifs have previously been substituted for alanine residues (WA and Gx3A) and such mutants are unable to support retrotranslocation (Fig. 5E & F) (Neal et al., 2018). In addition, we have previously identified that the L1 and TM2 regions of Dfm1 are critical for its retrotranslocation function (Fig. 5E & F). Specifically, L1 mutants (F58S, L64V, and K67E) impaired membrane substrate binding to Dfm1 and TM2 mutants (R98L, S99V, S100V, and Q101L) impaired Dfm1’s lipid thinning distortion function (Nejatfard et al., 2021). The lipid distortion function of Dfm1 increases lipid permeability to aid the extraction of integral membrane substrates from the lipid bilayer. The effect of these retrotranslocation-deficient mutants on Orm2 degradation was directly tested with the CHX-chase assay. All Dfm1 mutants, with the exception of Dfm1-5Ashp, completely stabilized Orm2 (Fig. 5G). Overall, the conserved rhomboid motifs, WR and Gx3G, the L1 region for substrate binding, and the TM2 region for lipid thinning, are all required for Orm2 degradation. We employed another functional assay for Dfm1 to test whether the retrotranslocation-deficient Dfm1 mutants can restore growth in *dfm1*Δ*orm1*Δ cells, which normally have impaired growth at 37°C due to increased flux in sphingolipid synthesis. To this end, adding back empty vector or wild-type DFM1 to *dfm1*Δ*orm1*Δ cells resulted in the expected impairment and rescue of normal growth, respectively (Fig. S2B & C). Introduction of Dfm1 mutants to *dfm1*Δ*orm1*Δ cells did not rescue growth defects, with the exception of Dfm1-5Ashp, which was able to rescue the growth defect in a manner similar to that of WT Dfm1 (Fig. S2B). Taken together, these data suggest that the substrate binding, lipid distortion function, and conserved rhomboid motifs, but not Cdc48 recruitment function, of Dfm1 are required for Orm2 degradation.

### Orm2 degradation is dependent on EGAD, but not ERAD or INMAD

Given that the Cdc48 recruitment function of Dfm1 is dispensable for Orm2 degradation, it seems likely that Dfm1’s retrotranslocation function in ERAD is not required for Orm2 degradation. Accordingly, we wished to survey for all protein degradation pathways in which Dfm1 may participate. The secretory pathway possesses several protein quality-control pathways including the INM-associated degradation (INMAD), ERAD, and EGAD, which govern both regulated and quality-control degradation of INM proteins, ER proteins, and Endosomal/Golgi proteins, respectively (Sicari et al., 2019; Sun and Brodsky, 2019). All pathways employ dedicated E3 ligases that determine substrate specificity and ubiquitination. Specifically, the Asi and Doa10 E3 ligases mediate INMAD, the Hrd1 and Doa10 E3 ligases mediate ERAD, and Tul1 E3 ligase mediates EGAD. A unifying theme for all protein degradation pathways is that they require the hexameric AAA ATPase, Cdc48, and the proteasome for retrotranslocation and degradation of all substrates. We utilized the CHX-chase assay to test the requirement for all E3 ligases, Cdc48, and the proteasome for degradation of Orm2. In line with previous studies (Schmidt et al., 2019), Orm2 was still degraded with similar kinetics to wild-type strains in *hrd1Δ, doa10Δ*, and *asi1Δ* (Fig. 6A). These results indicate that Orm2 degradation does not require either the INMAD or ERAD pathways. As expected, Orm2 degradation was completely inhibited in *tu1lΔ* cells, *cdc48-2* cells, and proteasome subunit mutant, *hrd2-1* (Fig. 6B) (Schmidt et al., 2019a). These observations are in accordance with previous studies and demonstrate that Orm2 ubiquitination, extraction, and proteasome degradation is mediated solely by EGAD (Schmidt et al., 2019). To further confirm that Orm2 degradation is independent of ERAD and INMAD, we performed *in vivo* ubiquitination assays on WT, *asi1Δ*, *hrd1Δ*, *doa10Δ*, *tul1Δ*, *cdc48-2*, and *hrd2-1* strains. Strains were lysed and subjected to immunoprecipitation (IP) using anti-RFP antibodies, followed by immunoblotting (IB) with anti-ubiquitin and anti-RFP antibodies. As suggested by CHX-chase experiments conducted by our lab and others (Schmidt et al., 2019), the degree of Orm2 ubiquitination in *asi1Δ*, *hrd1Δ*, and *doa10Δ* was similar to that seen in WT strains, demonstrating that the E3 ligases Asi, Hrd1, and Doa10 are not involved in the polyubiquitination of Orm2 (Fig. 6C, *lanes 1, 2, 3, 4*). In line with a previous study, the amount of Orm2 ubiquitination is increased in *cdc48-2* and *hrd2-1* cells, suggesting that Orm2 is on pathway for retrotranslocation and proteasome degradation (Fig. 6C, *lanes 8, 9*) (Schmidt et al., 2019).

**Fig. 6.**
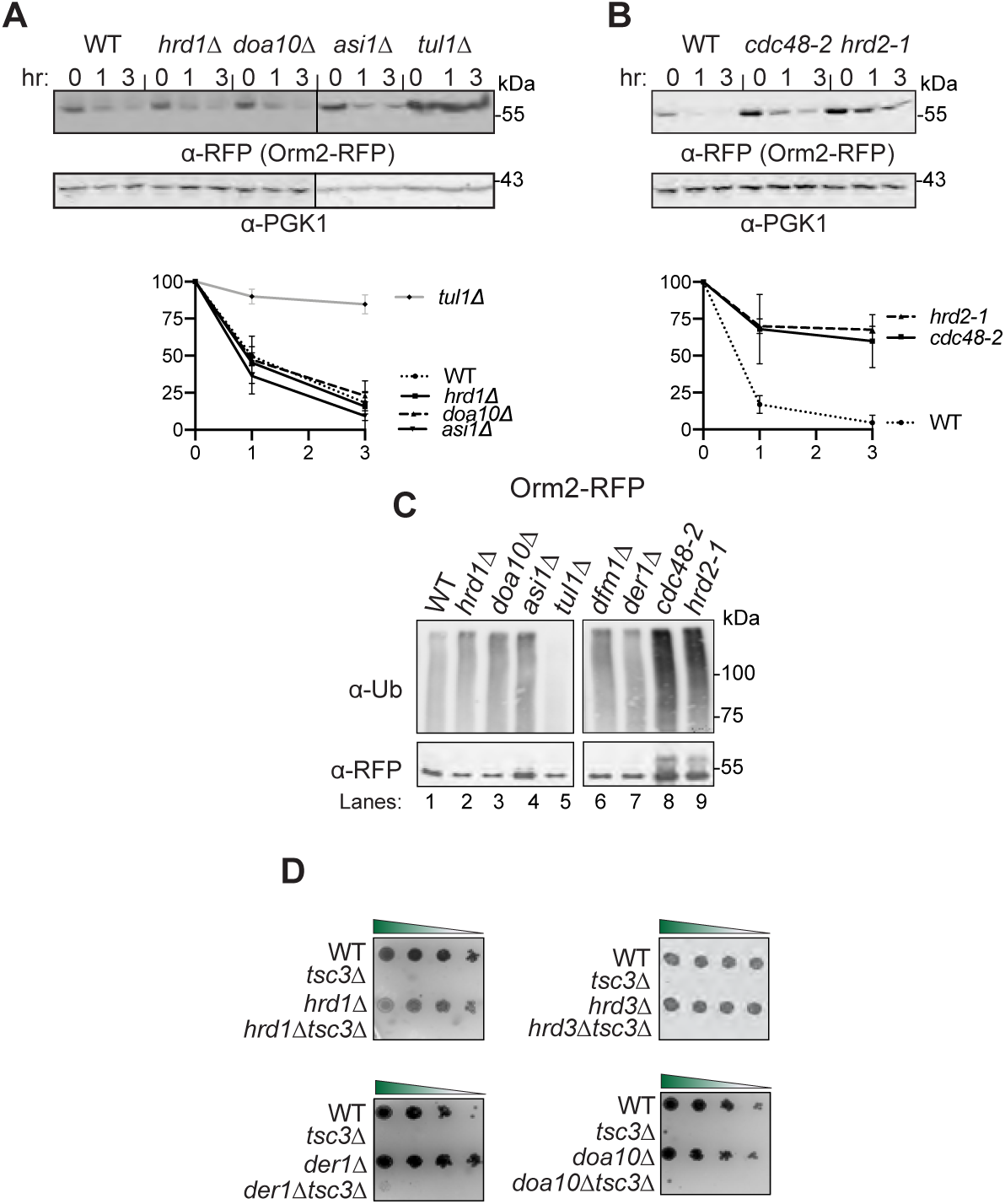
Orm2 degradation requires EGAD, but not ERAD or INMAD pathway. **(A)** E3 ligase Tul1 is required for Orm2 degradation. In the indicated strains, degradation of Orm2-RFP was measured by CHX-chase assay. Cells were analyzed by SDS-PAGE and immunoblotted for Orm2-RFP with α-RFP. **(B)** Cdc48 and the proteasome are required for Orm2 degradation. Same as (A), except *cdc48-2* and *hrd2-1* were analyzed for Orm2-RFP degradation. **(C)** Dfm1 does not function in the post-ubiquitination step of Orm2 degradation pathway. Indicated strains expressing Orm2-RFP were grown into log phase. Cells were lysed, and microsomes were collected and immunoprecipitated with α-RFP conjugated to agarose beads. Samples were then subjected to SDS-PAGE and immunoblot by α-Ubiquitin and α-RFP. One of three biological replicates is shown. **(D) ERAD mutants do not rescue temperature-sensitive lethality of *tsc3*Δ.** Indicated strains were grown to log-phase in SC and serially diluted cultures were plated on SC plates and incubated at 37°C and imaged at Day 2. One representative of three biological replicates is shown n=3.

As expected, Orm2 ubiquitination was decreased in *tul1Δ* strains, indicating that Orm2 is ubiquitinated by Tul1-dependent EGAD (Fig. 6C, lane 5). To further confirm that Orm2 degradation is independent of EGAD and INMAD, we next tested whether any of the ERAD components besides Dfm1, genetically interacted with Tsc3 and Orm1. Specifically, we examined whether any ERAD mutants phenocopy *dfm1Δtsc3Δ* cells, which rescues lethality at 37°C, or *dfm1Δorm1Δ*, which exhibits a growth defect at 37°C. To this end, double mutants were generated in which *tsc3Δ* or *orm1*Δ was knocked out, along with the following HRD and DOA pathway components: *hrd1Δ*, *hrd3*Δ, *der1*Δ, and *doa10*Δ. In all cases, the HRD and DOA pathway mutants did not phenocopy *dfm1*Δ: *hrd1Δtsc3Δ*, *hrd3Δtsc3Δ*, *der1Δtsc3Δ*, and *doa1Δtsc3Δ* were unable to rescue the temperature-sensitive lethality of *tsc3*Δ; and *hrd1Δorm1Δ, hrd3Δorm1Δ, der1Δorm1Δ,* and *doa10Δorm1Δ* did not exhibit an exacerbated growth defect at 37°C. Hence, the DOA and HRD ERAD pathways do not genetically interact with Tsc3 or Orm2. In summary, CHX-chase, genetics, and *in vivo* ubiquitination assays confirmed that Orm2 is degraded solely by the EGAD pathway and not by INMAD or ERAD.

### Dfm1 does not function at the post-ubiquitination step of Orm2 degradation pathway

To determine the step at which Dfm1 functions in Orm2 degradation, the ubiquitination status of Orm2 in *dfm1Δ* strains was analyzed. We have previously demonstrated that Dfm1 functions at the post-ubiquitination step of ERAD, with an increased degree of polyubiquitination of ERAD-M substrates was observed in *dfm1Δ* strains (Neal et al., 2018). This was caused by the inability of Dfm1 to retrotranslocate its substrates, resulting in build-up of polyubiquitinated membrane substrates along the ER membrane. Surprisingly, in *dfm1Δ* strains, the level of Orm2 ubiquitination was the same as in WT strains and did not phenocopy retrotranslocation-deficient strains, *cdc48-2*, or the proteasomal mutant *hrd2-1* (Fig. 6C, lanes 7,8,9). Hence, Dfm1 does not function in the post-ubiquitination step of EGAD. We also tested the requirement of Der1 for Orm2 ubiquitination and saw no change in Orm2 ubiquitination levels in *der1Δ* strains compared with WT strains (Fig. 6C, lanes 1 & 7). Taken together, these data suggest that Dfm1 does not function at the post-ubiquitination step of the Orm2 degradation pathway.

### Dfm1 does not directly function in EGAD

Given the requirement for Dfm1 in Orm2 degradation, it is surprising that the Dfm1-dependent ERAD pathway is not involved with Orm2 degradation. It is possible that Dfm1 directly functions in EGAD. To test this hypothesis, we examined the interaction of Dfm1 with the Dsc complex (E3 ligase Tul1 and Dsc2), which mediates substrate detection and ubiquitination within the Golgi in the EGAD pathway. Dfm1-GFP was immunoprecipitated with GFP Trap antibodies followed by SDS-PAGE and immunoblotting for endogenous Tul1 and Dsc2 with anti-Tul1 and anti-Dsc2, respectively. In all cases, no association of Dfm1 with Tul1 and Dsc2 was observed, while Dfm1 was able to associate with Cdc48, as expected (Fig. S3). Moreover, fluorescence microscopy demonstrated that Dfm1 is solely localized in the ER and does not co-localize with Golgi-associated markers (Fig. 2B). Finally, there were no significant interactions between Dfm1 and any EGAD components identified from our proteomic analysis (Fig. 1E).

### Dfm1 is required for Orm2 export from the ER to the Golgi

The observation that Dfm1-5Ashp can still facilitate Orm2 degradation suggest that Dfm1-mediated degradation of Orm2 is independent of its canonical retrotranslocation role in ERAD. Hence, we sought to identify the specific step at which Dfm1 functions within the Orm2 degradation pathway. EGAD-mediated degradation of Orm2 is most well characterized in yeast where it consists of five steps: 1) phosphorylation of Orm2 by Ypk1 in the ER, 2) COPII-mediated export of phosphorylated Orm2 from ER to Golgi and endosome, 3) polyubiquitination of Orm2 by the E3 ligase Dsc2, 4) retrotranslocation of substrates from the Golgi/endosome to the cytosol, and 5) degradation of the ubiquitinated substrates by the cytosolic proteasome (Schmidt et al., 2019). To determine which step was blocked in Dfm1-deficient cells, we analyzed the phosphorylation status of Orm2 in *dfm1*Δ cells. In *dfm1*Δ, Orm2 phosphorylation was increased to levels similar to those in the Dsc complex mutant *tul1*Δ, a knockout that blocks Orm2 degradation and leads to accumulation of phosphorylated Orm2 (Fig. 7A). This indicates *dfm1*Δ cells result in defective trafficking of Orm2 to the Golgi. To validate this in a cellular context, we next utilized live cell imaging fluorescence microscopy to determine the cellular compartment in which Orm2 was accumulating in *dfm1*Δ cells. In line with a previous study, Orm2 accumulated mainly in the early endosomes in *tul1*Δ cells (Schmidt et al., 2019), indicating that Orm2 was being routed to the Golgi/endosomes for degradation. By contrast, in *dfm1*Δ cells, Orm2 accumulated mainly at the ER and Orm2 co-localized with an ER, but not an endosome marker (Fig. 7B). In parallel, we utilized a phospho-mimetic Orm2 variant (Orm2-3D), which has been shown to mimic Ypk1-dependent constitutive phosphorylation and is continuously exported from the ER and degraded via EGAD. Indeed, we and others show that in WT cells, Orm2-3D is rapidly degraded (Fig. 7C) (Schmidt et al., 2019, 2020). Remarkably, by employing a CHX-chase assay, we found that Orm2-3D degradation was completely prevented in *dfm1*Δ cells (Fig. 7C). Using microscopy, we confirmed that Orm2-3D remained exclusively in the ER in *dfm1*Δ cells (Fig. 7E). By contrast, degradation of an Orm2 phosphonull variant (Orm2-3A) was completely prevented in WT cells (Fig. 7C) and we and others showed that Orm2-3A was not exported to the ER (Fig. S4)(Schmidt et al., 2019). Because *orm2*Δ cells can rescue *tsc3*Δ lethality, we wanted to test whether Orm2-3D or Orm2-3A elicits the same effect. To test this, either phosphonull Orm2-3A or phosphomimetic Orm2-3D were added to *tsc3*Δ*orm2*Δ cells and the growth assay was employed. As controls, Orm2-3A alone and Orm2-3D cells grew similarly as WT cells whereas *tsc3*Δ cells exhibited the expected growth lethality at 37°C. Notably, *tsc3*Δ*orm2*Δ cells containing Orm2-3D alleviated *tsc3*Δ lethality whereas *tsc3*Δ*orm2*Δ cells containing Orm2-3A did not rescue *tsc3*Δ lethality (Fig. 7D). These results suggest that two conditions are sufficient in rescuing *tsc3*Δ lethality: 1) absence of Orm2 (*orm2*Δ) or 2) continuous phosphorylation and degradation of Orm2 via EGAD (Orm2-3D). In summary, based on CHX-chase, live image florescence microscopy, and genetic interaction assays, we demonstrate that Dfm1 is required for the export of phosphorylated Orm2 from the ER to Golgi. The physiological consequence for Dfm1 dysfunction is retention of Orm2 in the ER, which prevents its subsequent degradation.

**Fig. 7.**
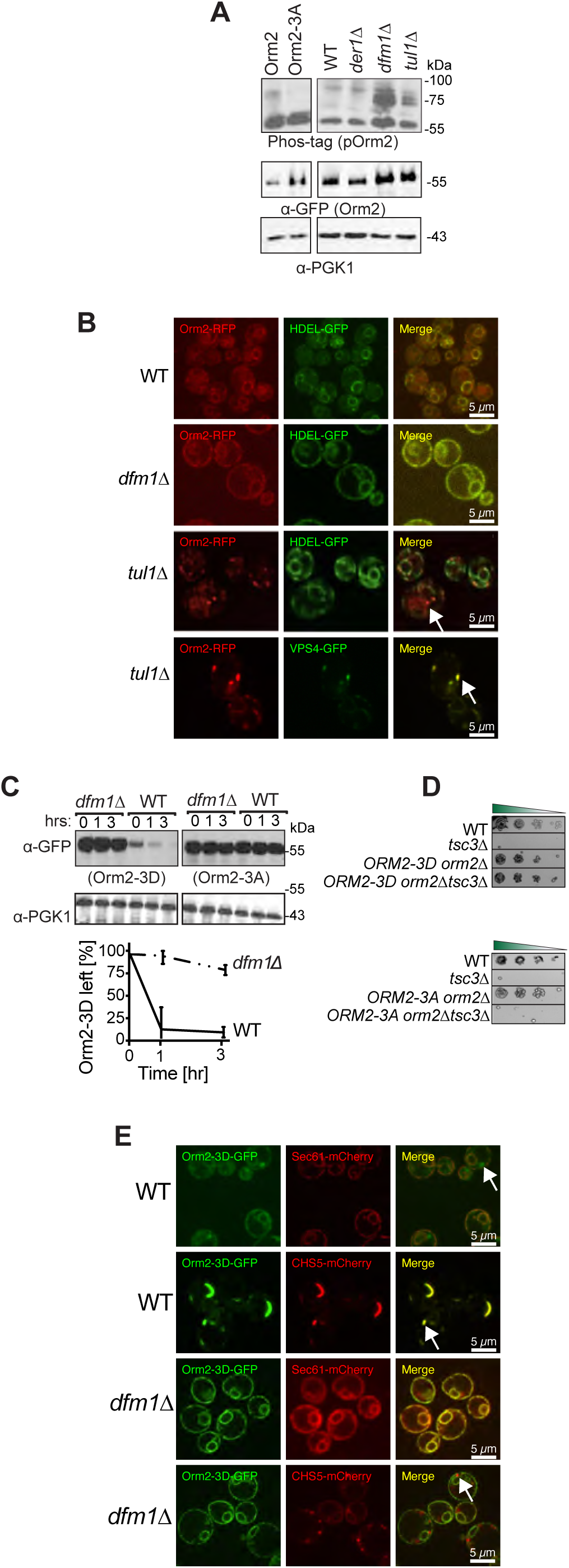
dfm1Δ accumulates phosphorylated Orm2 exclusively in the ER. **(A)** Phos-tag western blot analysis shows that there is an accumulation of phosphorylated Orm2 in *dfm1*Δ cells. The indicated strains were grown to log phase, treated with vehicle or 1.5 μM myriocin for 1 hour and subjected to SDS-PAGE or Phos-tag western blot analysis via blotting for Orm2 with α-RFP and PGK1 with α-PGK1 antibodies. **(B)** Orm2 is retained in the ER in *dfm1*Δ cells. Strains were grown to mid-exponential phase in minimal media and GFP and RFP fluorescence was examined on an AxioImager.M2 fluorescence microscope using a 100x objective and 28HE-GFP or 20HE-rhodamine filter sets (Zeiss). WT, *dfm1*Δ, and *tul1*Δ cells expressing Orm2-RFP. HDEL-GFP (ER marker, green) or VPS4-GFP (endosome marker, green) were used to test for co-localization with Orm2-RFP. *Scale bar, 5 μM*. *Arrowheads* indicate Orm2 co-localizing in post-ER compartments. **(C)** *dfm1*Δ cells block the degradation of phosphorylated mimic of Orm2 (Orm2-3D). In the indicated strains, degradation of Orm2-3A-GFP and Orm2-3D-GFP was measured by CHX-chase assay. Cells were analyzed by SDS-PAGE and immunoblotted α-GFP. **(D)** Accumulation of phosphorylated Orm2 within the ER is sufficient for rescuing the temperature-sensitive lethality of *tsc3*Δ cells. Indicated strains were spotted 5-fold dilutions on SC plates in triplicates, and plates were incubated at 37°C (n=3). **(E)** *dfm1*Δ blocks export of phosphorylated Orm2. Fluorescence imaging was performed as in (B) except WT and *dfm1*Δ cells expressing Orm2-3D-GFP was used. Sec61-mCherry (ER marker, red) or CHS5-mCherry (endosome marker, red) were used to test for co-localization with Orm2-3D-GFP. *Scale bar, 5 μM*. *Arrowheads* indicate Orm2 co-localizing in post-ER compartments.

### Loss of Dfm1 does not affect COPII-mediated trafficking

Because loss of Dfm1 resulted in accumulation of Orm2 in the ER, we directly interrogated the role of Dfm1 in COPII-mediated trafficking. To test whether Dfm1 has a direct function in COPII-mediated trafficking, we analyzed the steady-state levels of COPII cargo substrate, carboxypeptidase Y (CPY), and found that in *dfm1*Δ cells, the mature form of CPY accumulated at similar levels as WT cells (*m*; Fig. S4B). As a control for a deficiency in COPII-mediated export, when *sec12-1* cells were shifted to nonpermissive growth temperature at 37°C, there was the expected buildup of the premature form (*P*; Fig. S4B). Finally, we did not identify significant interactions between Dfm1 and any COPII trafficking components from our proteomic analysis (Fig. 1C). Taken together, our data suggests that Dfm1 does not directly function in COPII-dependent trafficking.

## DISCUSSION

In this study, we describe a novel role for the derlin rhomboid pseudoprotease, Dfm1, in maintaining sphingolipid homeostasis. The function of Dfm1 in ERAD-M retrotranslocation of misfolded membrane protein substrates has been well established in our laboratory and this study uncovers an additional biological function of Dfm1. The role of Dfm1 in maintaining sphingolipid homeostasis appears to be separate from its role in ERAD-M retrotranslocation. Specifically, our studies indicate that Dfm1 is required to facilitate the export of phosphorylated Orm2 from the ER and that this function requires Dfm1’s substrate binding and lipid thinning activity, but not its Cdc48 recruitment function. Overall, our studies reveal a novel role for rhomboid pseudoproteases in maintaining sphingolipid homeostasis, a function that is independent of their role in ERAD.

To identify Dfm1 interacting partner proteins, we performed proximity-dependent biotinylation (BioID) coupled with mass spectrometry. Several proteins found in close proximity to Dfm1 were involved in the sphingolipid biosynthesis pathway. The first committed step of the sphingolipid biosynthetic pathway is catalyzed by the serine palmitoyl transferase (SPT) complex, which consists of Lcb1, Lcb2, and Tsc3. This step is strictly regulated by Orm1/Orm2 and Sac1, which negatively regulates SPT, and Tsc3, and enhances SPT activity 100-fold. Indeed, we confirmed a physical interaction between Dfm1 and the SPOTS complex members Orm2 and Lcb1. To further explore the relationship between Dfm1 and SPOTS complex members, we examined the genetic interactions between *dfm1*Δ and knockout of SPOTS complex members or sphingolipid biosynthetic enzymes. We exploited the *tsc3*Δ growth lethality phenotype at 37°C, which is due to Tsc3 being required for enhancing SPT activity at 37°C. Growth assays and lipidomic analyses indicated that *dfm1*Δ cells phenocopy established negative regulators of the SPT enzymes, *orm1*Δ and *orm1*Δ, where all three are able to reverse the temperature-sensitive lethality of *tsc3*Δ by increasing ceramide and complex sphingolipid levels. Therefore, we propose that Dfm1 antagonizes the sphingolipid biosynthesis pathway.

The mechanism associated with Dfm1-dependent Orm2 export from the ER remains to be elucidated. Our data suggests that Dfm1 functions at the post-phosphorylation step of Orm2 where loss of Dfm1 blocks ER export of phosphorylated Orm2. Moreover, Dfm1 does not directly function in COPII export since its absence does not abrogate export of a COPII cargo, CPY. Based on these findings, Dfm1 most likely functions upstream of COPII-dependent trafficking. However, we cannot exclude the possibility that Dfm1 may directly participate in Orm2 export in a COPII-independent manner. The former model seems plausible since our laboratory has recently identified a chaperone-like Dfm1 function that is required for disaggregating misfolded membrane substrates along the ER membrane (Kandel et al., 2022). Similar to Dfm1’s function as a mediator in sphingolipid homeostasis, its chaperone-like role requires Dfm1’s substrate and lipid thinning function, but not its Cdc48 recruitment function. Accordingly, we surmise that Dfm1 utilizes its chaperone-like activity to target phosphorylated Orm2 within the SPOTS complex and deliver it to the COPII machinery for ER export (Fig. 8). The extent to which the chaperone-like activity of Dfm1 is required for Orm2 export from the ER warrants future investigations.

**Fig. 8.**
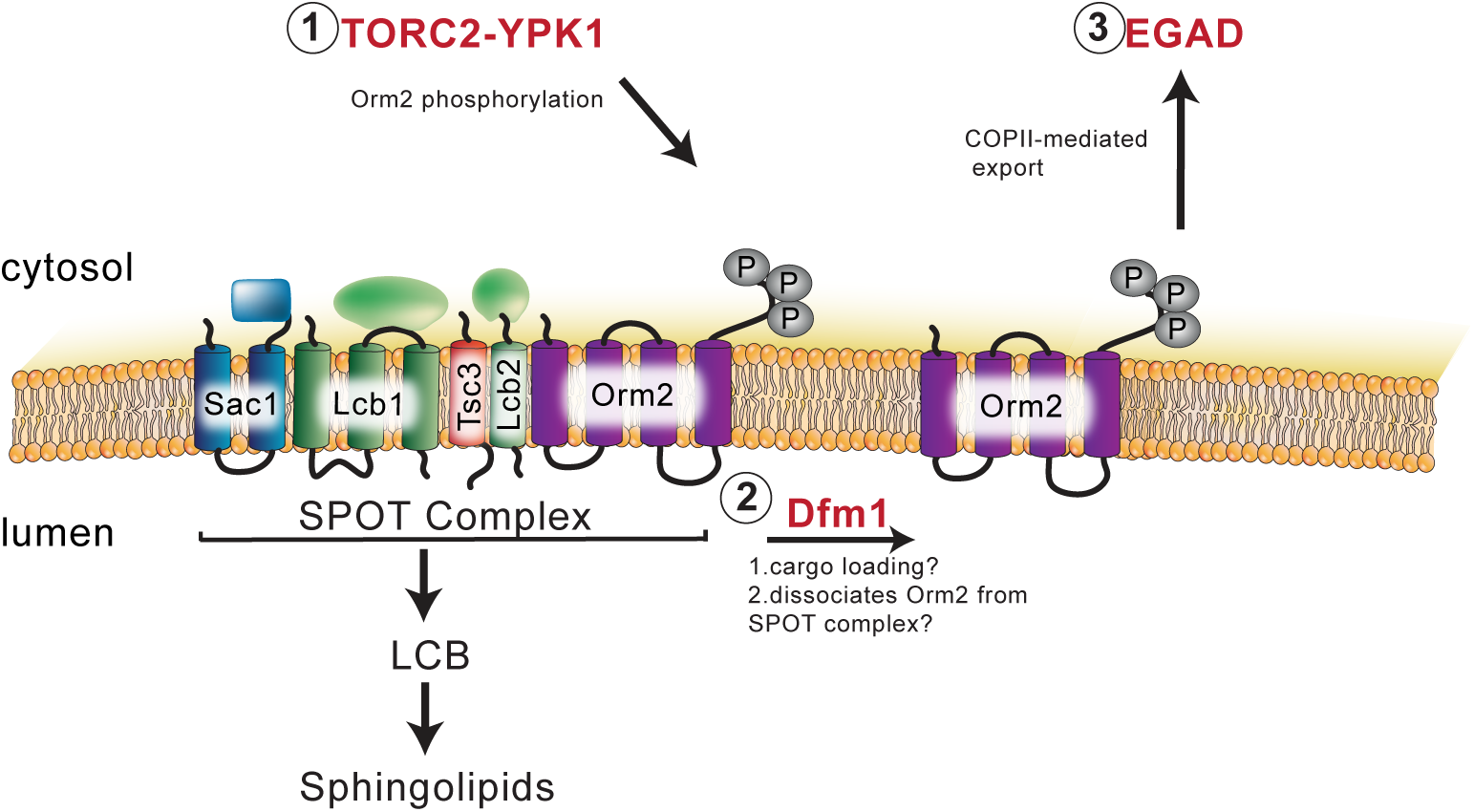
Schematic of Dfm1’s role in Orm2 degradation. 1) Orm2 is inactivated via phosphorylation by TORC2-YPK1 signaling axis. 2) Dfm1 functions downstream of Orm2 phosphorylation and possibly plays a role in delivering phosphorylated Orm2 to COPII vesicles. 3) Phosphorylated Orm2 is routed to the Golgi and degraded via EGAD.

Our data demonstrates that Dfm1-deficient cells are accumulating Orm2, the negative regulator of sphingolipid biosynthesis, which raises the question of why increased levels of LCBs, ceramides, and complex sphingolipids were observed in *dfm1*Δ and *dfm1*Δ*tsc3*Δ cells? Our results were in contrast to our initial expectation that accumulation of Orm2 would decrease levels of LCB and ceramides, since Orm2 antagonizes the sphingolipid biosynthesis pathway. One possibility is that in *dfm1*Δ cells, the majority of Orm2 in the ER is in a phosphorylated and inactive state –as shown in Fig. 7A–and is no longer able to repress SPT activity. This is supported by a previous study in which Orms have been found to directly regulate the localization and oligomerization state of SPT at the ER in a manner that is dependent upon their phosphorylation state. Specifically, phosphorylated Orm2 shifts SPT to the monomeric state and contributes to sustained SPT activity (Han et al., 2019). This suggests that ER-localized phosphorylated Orm2 is no longer active, supporting our observation in which LCB levels are increased in *dfm1*Δ cells.

The ER hosts metabolic pathways that synthesize a variety of lipids such as phospholipids, cholesterol, and sphingolipids. Thus, it is critical for the ER to sense and respond to fluctuations in lipid composition in order to maintain cellular homeostasis (Fun and Thibault, 2020; Piña et al., 2018; Tam et al., 2018). Several pathways are employed to maintain the flux of the lipid biosynthetic pathway. One such pathway is the targeting and degradation of lipid biosynthetic enzymes through ERAD-mediated degradation. For example, cholesterol synthesis is downregulated through regulated ERAD of the rate-controlling cholesterol biosynthetic enzymes Hmg-CoA reductase and squalene monooxygenase (Wangeline et al., 2017). Maintenance of lipid homeostasis is also critical for protein synthesis, as dysregulated lipid levels negatively impact ER protein quality-control machineries. This is supported by studies demonstrating that increased flux of sphingolipids induces UPR and sensitivity of cells to ER stress (Han et al.). Previous studies from our lab and others have demonstrated that lipid thinning by the ERAD machinery facilitates the removal of misfolded substrates from the ER (Neal et al., 2018; Wu et al., 2020). Because lipids play a large role in ERAD retrotranslocation, fine-tuning lipid levels is critical to ensure that ERAD remains intact and functional. Our findings implicate Dfm1 as a mediator in sphingolipid homeostasis. Interestingly, by utilizing homology modeling and bionformatic analysis, we identified sphingolipid-binding motifs on TM1 and TM6 of Dfm1, suggesting that Dfm1 may directly detect sphingolipid levels and fine-tune the control of sphingolipid production by modulating the export of Orm2 (unpublished data). Similarly, cholesterol has been shown to directly regulate levels of the E3 ligase MARCH6 levels, with increased MARCH6 levels leading to ubiquitination and degradation of the key cholesterol enzyme, squalene monoozygenase (Zelcer et al., 2014). Future studies on the lipid-sensing function of Dfm1 and how it de-represses SPT activity via Orm2 export from the ER will require additional experimentation.

The Orm family proteins are well conserved and all three human ORMDLs associate with SPT and directly regulate SPT activity. However, unlike their yeast counterparts, they do not appear to be phosphorylated since they lack the homologous N-terminal domains that are phosphorylated by Npr1, Ypk1, and Ypk2 in yeast. Instead, ORMDL protein levels are regulated directly by ceramide levels (Davis et al., 2019). Consistent with this finding, altered protein levels of ORMDLs are associated with the pathophysiology of a range of diseases, including colorectal cancer, inflammation, obesity, and diabetes (Davis et al., 2018). Moreover, single nucleotide polymorphisms near ORMDL3, which lead to its increased expression, are associated with childhood asthma. Accordingly, our observation that yeast Dfm1 alters Orm2 protein levels and impacts sphingolipid metabolism raises the possibility that derlins have a causative role in these diseases. Defining the mechanism of Dfm1-mediated regulation of Orm2 levels should illuminate new treatment paradigms for patients with dysregulated sphingolipid metabolism.

In summary, we have performed proteomic analyses to enable the identification of Dfm1 interacting factors. These studies have demonstrated that several key regulators of sphingolipid biosynthesis are associated with Dfm1 and we report a novel function for Dfm1 in mediating sphingolipid homeostasis. Sphingolipids play diverse roles in cellular functions, which includes cell signaling, supporting cellular structure, providing energy storage, and regulating cell growth cycles. Dysregulation of sphingolipid levels has been associated with several life-threatening disorders. Overall, this study identifies derlin rhomboid pseudoproteases as key regulators of sphingolipid levels and reveals them as potential therapeutic targets for treatment of lipid disorders that are associated with the dysregulation of sphingolipid levels.

## AUTHOR CONTRIBUTIONS

S.B., A.A., Y.O., M.P., R.K., E.B., A.K., and S.E.N. designed research; S.B., A.A., Y.O., M.P., J.J., I.I., R.K., I.W., N.S., and A.F. performed research; S.B., A.A., Y.O., M.P., J.J., I.I., R.K., I.W., N.S., and A.F analyzed data; S.B., A.A., and S.E.N. wrote the paper; and S.B. and S.E.N. designed illustrations for figures. All authors reviewed the results and approved the final version of the manuscript.

## ACKNOWLEDGEMENTS

We thank Peter Espenshade (Johns Hopkins Medicine), Oliver Schmidt (Medical University of Innsbruck), David Teis (Medical University of Innsbruck), Teresa Dunn (National Institutes of Health), and Peter Novick (University of California, San Diego) for providing plasmids, yeast strains, antibodies. We also thank Dr. Maho Niwa, Dr. David Teis, Dr. Oliver Schmidt, and the Neal lab members for in depth discussions and technical assistance. These studies were supported by NIH grant 1R35GM133565-01, Pew Biomedical Award, and NSF CAREER grant (to S.E.N), HHMI Gilliam Fellowship GT15096 (to S.E.N and A.A.), and KAKENHI grant JPSSH04986 (to A.K.).

## RESOURCE AVAILABILITY

### Lead contact

Further information and requests for resources and reagents should be directed to and will be fulfilled by the Lead Contact, Sonya Neal.

### Materials availability

Plasmids and yeast strains generated in this study are available from our laboratory.

### Data and Code Availability

This study did not generate/analyze [dataset/code].

## METHOD DETAILS

### Yeast and Bacteria Growth Media

Standard yeast *Saccharomyces cerevisiae* growth media were used as previously described (Hampton and Rine, 1994), including yeast extract-peptone-dextrose (YPD) medium and ammonia-based synthetic complete dextrose (SC) and ammonia-based synthetic minimal dextrose (SD) medium supplemented with 2% dextrose and amino acids to enable growth of auxotrophic strains at 30°C. *Escherichia coli* Top10 cells were grown in standard LB media with ampicillin at 37°C as previously described (Gardner et al., 1998). HEK293 cells were cultured in DMEM medium supplemented with 10% FBS.

### Plasmids and Strains

Plasmids used in this study are listed in Table S1. Plasmids for this work were generated using standard molecular biological techniques (Sato et al., 2009) and verified by sequencing (Eton Bioscience, Inc.). Primer information is available upon request. Lcb1-RFP and Orm2-RFP plasmids were a gift from Theresa Dun (Uniformed Services University of the Health Sciences, MD). Orm2-3A-GFP and Orm2-3D-GFP plasmids were a gift from Oliver Schmidt and David Teis (Medical University of Innsbruck, Austria).

A complete list of yeast strains and their corresponding genotypes are listed in Table S2. All strains used in this work were derived from S288C or Resgen. Yeast strains were transformed with DNA or PCR fragments using the standard LiOAc method (Ito et al., 1983). Null alleles were generated by using PCR to amplify a selection marker flanked by 50 base pairs of the 5’ and 3’ regions, which are immediately adjacent to the coding region of the gene to be deleted.

The selectable markers used for making null alleles were genes encoding resistance to G418 or CloNat/nourseothricin. After transformation, strains with drug markers were plated onto YPD followed by replica-plating onto YPD plates containing (500 μg/mL G418 or 200 μg/mL nourseothricin). All gene deletions were confirmed by PCR.

### *dfm1*Δ strain handling

Due to rapid suppression nature of *dfm1*Δ null strains, freshly transformed *dfm1*Δ null cells with the respective substrates should be used in every assay. Generation of Dfm1 mutant strains and troubleshooting guidelines are found in (Bhaduri and Neal, 2021).

### Proximity dependent-biotinylation (BioID) coupled with mass spectrometry

E.coli biotin ligase, BirA-3x Flag (pMP780) and Dfm1 (pSN113) were cloned under the Gal-2µM promoter contained in the vector pSN5. BirA was also cloned with wild-type SEN1 cells to be used as control. Both WT-BirA and Dfm1-BirA cells were inoculated into YNB-biotin media, supplemented with 1µM biotin, amino acids and 0.2% raffinose, and grown overnight. The following day, the cells were dialed back to 0.2OD and grown for 4 hours before addition of 0.2% galactose to induce the expression of BirA and Dfm1-BirA, incubated for 30 min, harvested and stored at −80°C overnight. The next day the cells were thawed and lysed using liquid nitrogen and incubated with preactivated Dynabeads (ThermoFisher) for 4 hours. The beads were subsequently seperated from the flowthrough using a magnetic stand, washed five times with PBS to separate the non-biotinylated proteins. Protein concentration was measured using a nanodrop, and samples were analyzed by mass-spectrometry to analyze the biotinylated interaction partners of Dfm1as previously described (Zuzow et al., 2018).

### Spot dilution growth assay

Yeast strains were grown in YPD or minimal selection media (-Leu -Ura) supplemented with 2% dextrose to log phase (OD_600_ 0.2-0.3) at 30°C. 0.2 OD cells were pelleted and resuspended in 500 μL dH_2_O. 12 μL of each sample was transferred to a 96-well plate where a five-fold serial dilution in dH_2_O of each sample was performed to obtain a gradient of 0.2-0.0000128 OD cells. The 8×12 pinning apparatus was used to pin cells onto synthetic complete (-Ura) agar plates supplemented with 2% dextrose or 2% galactose. Droplets of cells were air-dried in sterile conditions, then the plates were sealed with parafilm and incubated at 30°C. Plates were removed from the incubator for imaging after 3 days and again after 7 days.

### Cycloheximide-Chase Assay

Cycloheximide chase assays were performed as previously described (Sato et al., 2009). Cells were grown to log-phase (OD_600_ 0.2-.03) and cycloheximide was added to a final concentration of 50 μg/mL. At each time point, a constant volume of culture was removed and lysed. Lysis was initiated with addition of 100 μl SUME with protease inhibitors (PIs) and glass beads, followed by vortexing for 4 min. 100 μl of 2xUSB was added followed by incubation at 55°C for 10 min. Samples were clarified by centrifugation and analyzed by SDS-PAGE and immunoblotting.

### Fluorescence Microscopy

To prepare cells, overnight cultures were diluted to ∼0.20 OD in minimal media. After growing ∼3 hours, to log-phase (OD_600_∼.3-.6) samples were pelleted and washed with dH2O.

Fluorescence microscopy was accomplished using a CSU-X1 Spinning Disk (Yokogawa) confocal microscope at the Nikon Imaging Center on the UCSD campus.

### Native Co-IP

Cultures from various yeast strains were grown to OD_600_ .2-.45 and 15 ODs of cells were pelleted, rinsed with H_2_0 and lysed with 0.5 mM glass beads in 400 μL of MF buffer supplemented with protease inhibitors. This was followed by vortexing at 1-minute intervals for 6-8 minutes at 4°C. Lysates were combined and clarified by centrifugation at 2,500 g for 5 min followed by centrifugation at 14,000 g for 15 min to obtain the microsomal pellet. The microsomal pellet was resuspended in 1 mL of Tween IP buffer (500 mM NaCl, 50 mM Tris, pH 7.5, 10 mM EDTA. 1.5% Tween-20) and incubated on ice for 30 minutes. Lysates were then centrifuged for 30 min at 14,000 x g, and the supernatant was incubated overnight with 10 μL of equilibrated GFP-Trap® agarose (ChromoTek Inc., Hauppauge, NY) at 4°C. The next day, the GFP-Trap® agarose beads were combined to one tube, washed once with non-detergent IP buffer, washed once more with IP wash buffer and resuspended in 100 μL of 2xUSB. Samples were resolved on 8% SDS-PAGE and immunoblotted for Lcb1-RFP and Orm2-RFP α-RFP and Dfm1-GFP with α-GFP.

### *In vivo* ubiquitination assay

Western blotting to detect in vivo ubiquitination was performed as described previously (Garza et al., 2009). Briefly, yeast strains were grown to log phase (OD600 of 0.2 to 0.3). 15 OD equivalents of cells were pelleted by centrifugation and resuspended in lysis buffer (0.24 M sorbitol, 1 mM EDTA, 20 mM KH2PO4, pH 7.5) with PIs, after which 0.5 mm glass beads were added to the meniscus. The cells were lysed by vortexing in 1-min cycles at 4° C, with 1 min on ice in between, for 6 to 8 cycles. Lysates were clarified by centrifugation at 2,500 x g for 5 min. The clarified lysates were moved to fresh tubes, and 600 µL immunoprecipitation buffer (IPB; 15mM Na2HPO4, 150mM NaCl, 2% Triton X-100, 0.1% SDS, 0.5% deoxycholate, 10mM EDTA, pH 7.5) and 20 µL of GFP Trap (Chromotek) were added. Samples were incubated on ice for 5 min, clarified by centrifugation at 14,000 x g for 5 min, and moved to a fresh tube. Tubes were incubated at 4° C overnight with rocking. Beads were washed twice with IPB and then washed once with IP wash buffer (50 mM NaCl, 10 mM Tris, pH 7.5). Beads were aspirated to dryness, resuspended in 55 µL 2x USB, and incubated at 65° C for 10 minutes. Samples were resolved by SDS-PAGE on 10% gels, transferred to nitrocellulose, and immunoblotted with monoclonal anti-ubiquitin (Fred Hutchinson Cancer Research Institute) and anti-RFP (ThermoFisher) primary antibodies followed by goat anti-mouse (Jackson ImmunoResearch Laboratories) or goat anti-rabbit (Bio-Rad) HRP conjugated secondary antibody.

### Lipid analyses

Yeast cells (1 *A*_600_ units) were suspended in 100 μL of extraction solution [ethanol/water/diethyl ether/pyridine/15 M ammonia (15:15:5:1:0.018, v/v)], mixed with internal standards, and incubated at 60 °C for 15 min. As internal standards, four types of ceramides containing nine deuterium atoms (*d*_9_) [*N*-palmitoyl(*d*_9_)-dihydrosphingosine (*d*_9_-C16:0 Cer-A), *N*-palmitoyl(*d*_9_)-D-*ribo*-phytosphingosine (*d*_9_-C16:0 Cer-B), *N*-(2’-(*R*)-hydroxypalmitoyl(*d*_9_))-D-*erythro*-sphinganine (*d*_9_-C16:0 Cer-B’), *N*-(2’-(*R*)-hydroxypalmitoyl(*d*_9_))-D-*ribo*-phytosphingosine (*d*_9_-C16:0 Cer-C) (all purchased from Avanti Polar Lipids, Alabaster, AL)] were used (5 pmol each). After centrifugation (2,300 × *g*, room temperature, 2 min), the supernatant was recovered, and the pellets were suspended in 100 μL of extraction solution and incubated at 60 °C for 15 min again. After centrifugation (2,300 × *g*, room temperature, 2 min), the supernatant was recovered. The two supernatants were pooled, mixed with 700 µL of chloroform/methanol (1:2, v/v). To hydrolyze glycerolipids, alkaline treatment was performed by adding 37.5 µL of 3 M KOH and incubating at 37 °C for 30 min. After neutralization by adding 22.5 μL of 5 M formic acid, the samples were sequentially mixed with 250 µL of chloroform and 250 µL of water and centrifuged (20,400 × *g*, room temperature, 3 min) for phase separation. The organic phase containing lipids was recovered and dried. Lipids were resuspended in 625 μL of chloroform/methanol/water (5:4:1, v/v) and subjected to liquid chromatography (LC)-coupled tandem mass spectrometry (MS/MS) using a triple quadrupole mass spectrometer Xevo TQ-S (Waters, Milford, MA, USA) via multiple reaction monitoring and positive ion modes. The settings for LC separation and electrospray ionization were as described previously (Ohno et al., 2017)and the used *m/z* values and collision energies in the MS/MS measurement were listed in Supplementary Table 3. Ceramides were quantified by calculating the ratio of the peak area of each ceramide species to that of the internal standard corresponding to each type of ceramides. D-type ceramides were quantified using C-type ceramide standard (*d*_9_-C16:0 Cer-C).

## QUANTIFICATION AND STATISTICAL ANALYSIS

ImageJ (NIH) was used for all western blot quantifications. “Mean gray value” was set for band intensity measurements. In such experiments, a representative western blot was shown and band intensities were normalized to PGK1 loading control and quantified. t=0 was taken as 100% and data is represented as mean ± SEM from at least three experiments. GraphPad Prism was used for statistical analysis. Nested t-test, unpaired t-test or one-way factorial ANOVA followed by Bonferroni’s post-hoc analysis was applied to compare data. Significance was indicated as follow: n.s, not significant; * p<0.05, ** p<0.01, *** p<0.001, **** p<0.0001. The investigators were blinded during data analysis.

---

**Fig. S1.**
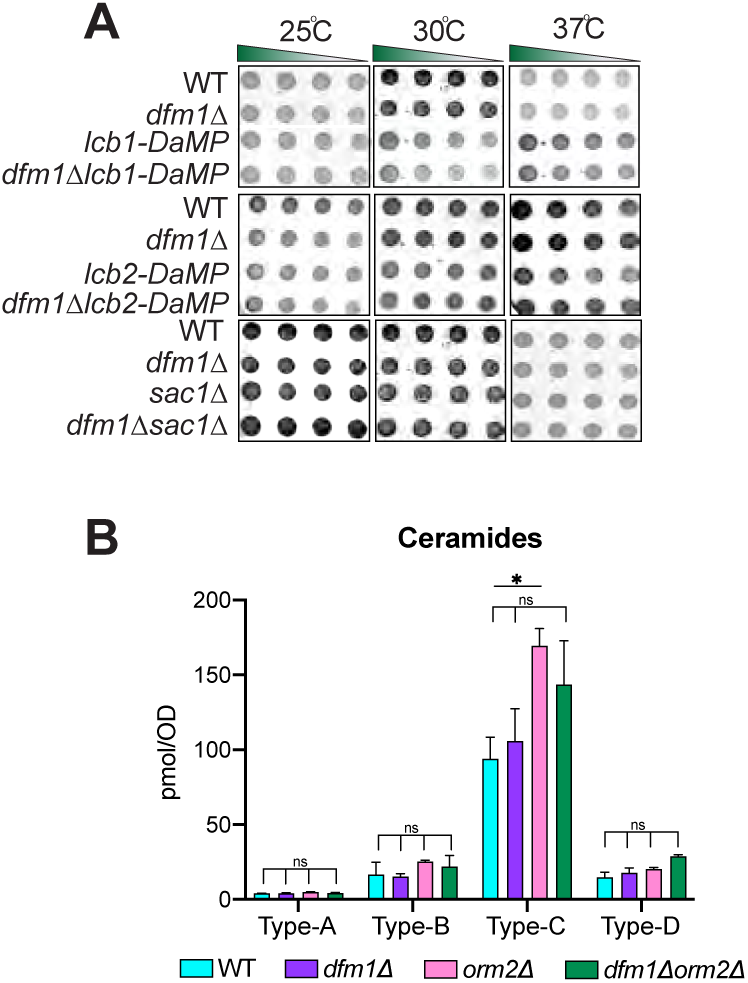
**(A)** Dfm1 doesn’t genetically interact with Lcb1, Lcb2, and Sac1. Indicated strains were spotted 5-fold dilutions on SC plates in triplicates, and plates were incubated at room temperature, 30°C, and 37°C (n=3). *Upper panel:* WT, *dfm1*Δ, *Lcb1-DaMP, and dfm1*Δ*Lcb1*-DaMP were compared for growth by dilution assay. *Middle panel:* WT, *dfm1*Δ, *Lcb2-DaMP, and dfm1*Δ*Lcb2*-DaMP were compared for growth by dilution assay. *Bottom panel:* WT, *dfm1*Δ, *and dfm1*Δ *sac1*Δ, were compared for growth by dilution assay. **(B)** WT, *dfm1*Δ, *orm2*Δ, and *orm2*Δ*dfm1*Δ cells were grown to log-phase at 30°C and lipids were extracted and subjected to LC-MS/MS. A- B- C- & D-Type ceramides containing C16, C18, C20, C22, C24, and C26 fatty acids were measured. Values represent the means±s.d.s of three independent experiments. Statistically significant differences compared to WT cells are indicated (Dunnett’s test; ns=non-significant).

**Fig. S2.**
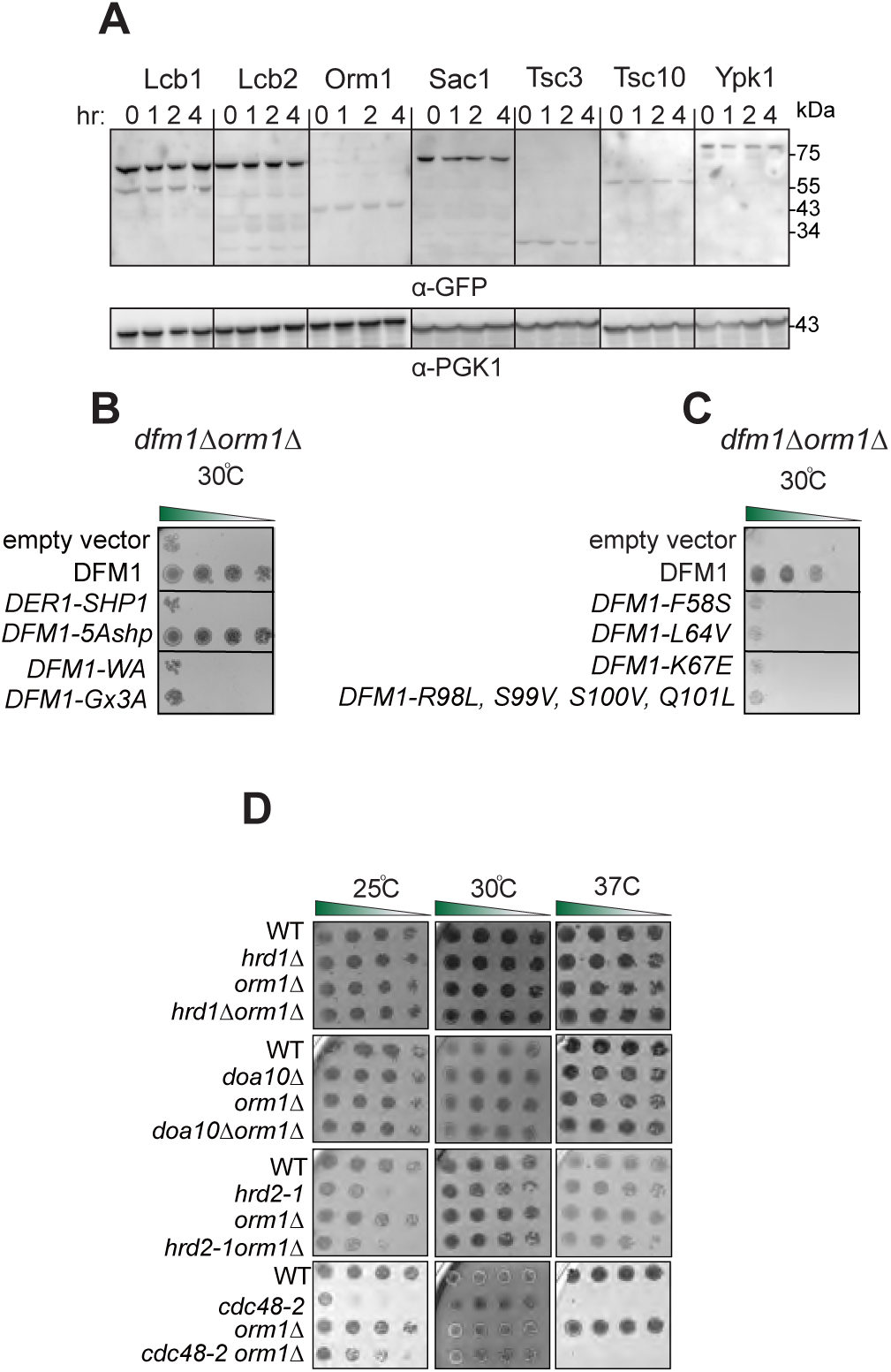
**(A)** Orm2 is degraded in WT strains. In the indicated WT strains, degradation of Lcb1-GFP, Lcb2-GFP, Orm2-GFP, Orm2-GFP, Sac1-GFP, Tsc3-GFP, Tsc10-GFP, and Ypk1-GFP was measured by CHX-chase assay. Cells were analyzed by SDS-PAGE and immunoblotted with α-GFP. **(B)** Serial dilution growth assay was performed on *dfm1*Δ*orm1*Δ and strains with DFM1, DER1-SHP, DFM1-AA, DFM1-Ax3A, DFM1-5Ashpmtnt, and empty vector addback. **(C)** Same as (B), except serial dilution growth assay was performed on *dfm1*Δ*orm1*Δ strains with L1 mutant addback: F58S, L64V, K67E and TMD2 quad mutant addback: DFM1-R98L, S99V, S100V, Q101L. Indicated strains were grown on SC-Leu plates at room temperature, 30°C and 37°C, and imaged on Day 2 and Day7. **(D)** ERAD mutants do not genetically interact with *orm1*Δ. Indicated strains were spotted 5-fold dilutions on SC plates in triplicates, and plates were incubated at room temperature, 30°C, and 37°C (n=3).

**Fig. S3.**
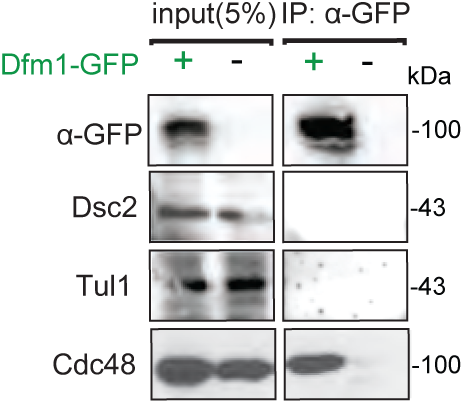
Dfm1 does not interact with EGAD components. Dfm1-GFP binding to EGAD members, Dsc2 and Tul1, were analyzed by co-IP. As negative control, cells not expressing Dfm1-GFP were used.

**Fig. S4.**
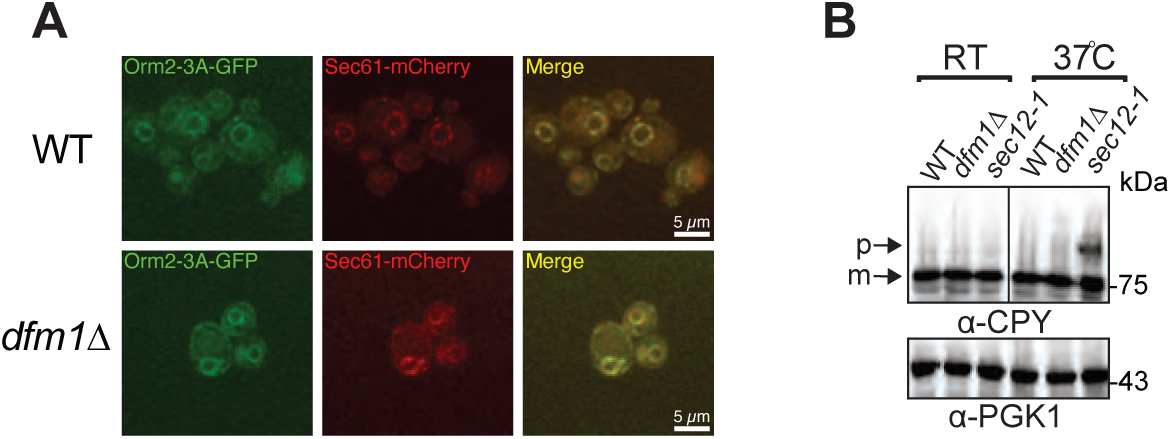
**(A)** Orm2-3A accumulates exclusively in the ER. Fluorescence imaging was performed as in Fig. 7B except WT and *dfm1*Δ cells expressing Orm2-3A-GFP was used. Sec61-RFP (ER marker, red) was used to test for co-localization with Orm2-3A-GFP. *Scale bar, 5 μM*. *Arrowheads* indicate Orm2 co-localizing in post-ER compartments. **(B)** *dfm1*Δ cells do not abrogate COPII-mediated export of CPY. The indicated cells were either grown at room temperature or shifted to non-permissive growth at 37°C. Cells were analyzed by SDS-PAGE and immunoblotted for CPY with α-CPY and PGK1 with α-PGK1.

**Table S1.** Plasmids used in this study, Related to Figures 1–7

| Plasmid | Gene |  |
| --- | --- | --- |
| pSN246 | YCp URA3 | pORM2-RFP |
| pSN247 | YCp LEU2 | pLCB1-RFP |
| pSN266 | YCp URA3 | pORM2-GFP |
| pSN267 | YCp URA3 | pORM2-3D-GFP |
| pSN268 | YCp URA3 | pORM2-3A-GFP |
| pSN60 | YCp LEU2 | pDFM1-L64V |
| pSN90 | YCp LEU2 | pDFM1 |
| pSN93 | YCp LEU2 | pDFM1-K67E |
| pSN230 | YCp LEU2 | pDFM1-R98L, S99A, S100A, Q101L |
| pSN95 | YCp LEU2 | pDFM1-F58S |
| pSN162 | YCp LEU2 | pDFM1-5Ashp-3HA |
| pSN161 | YCp LEU2 | pDER1-SHP-3HA |
| pSN164 | YCp LEU2 | pDFM1-AR-3HA |
| pSN163 | YCp LEU2 | pDFM1-Ax<sub>3</sub>G-3HA |
| pSN269 | YCp TRP1 | pORM2-RFP |
| pSN261 | YCp TRP1 | CHS5-mCherry |
| pSN258 | YCp LEU2/HIS3 | SEC61-mCherry |
| pSN252 | YCp URA3 | HDEL-GFP |

**Table S2.**
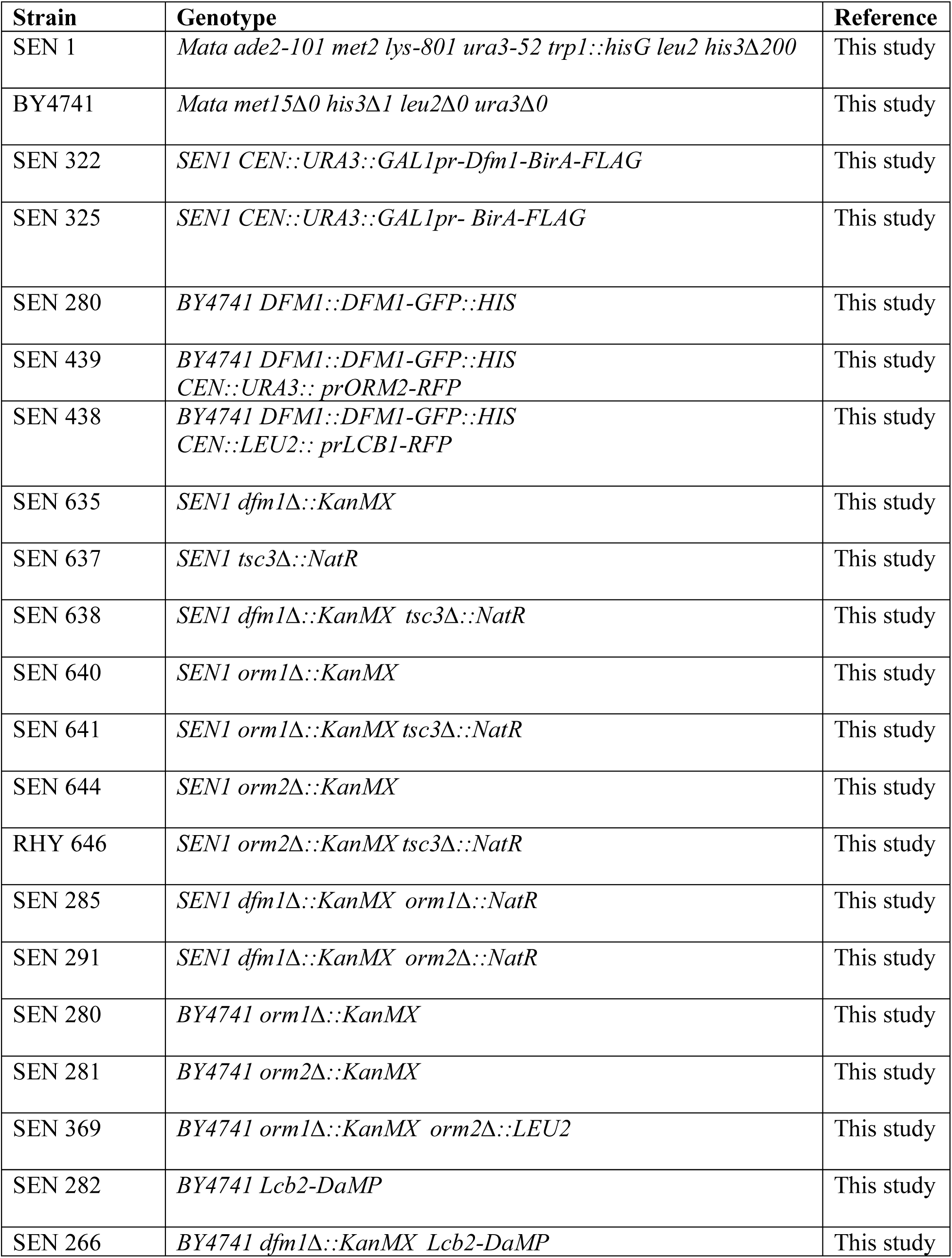
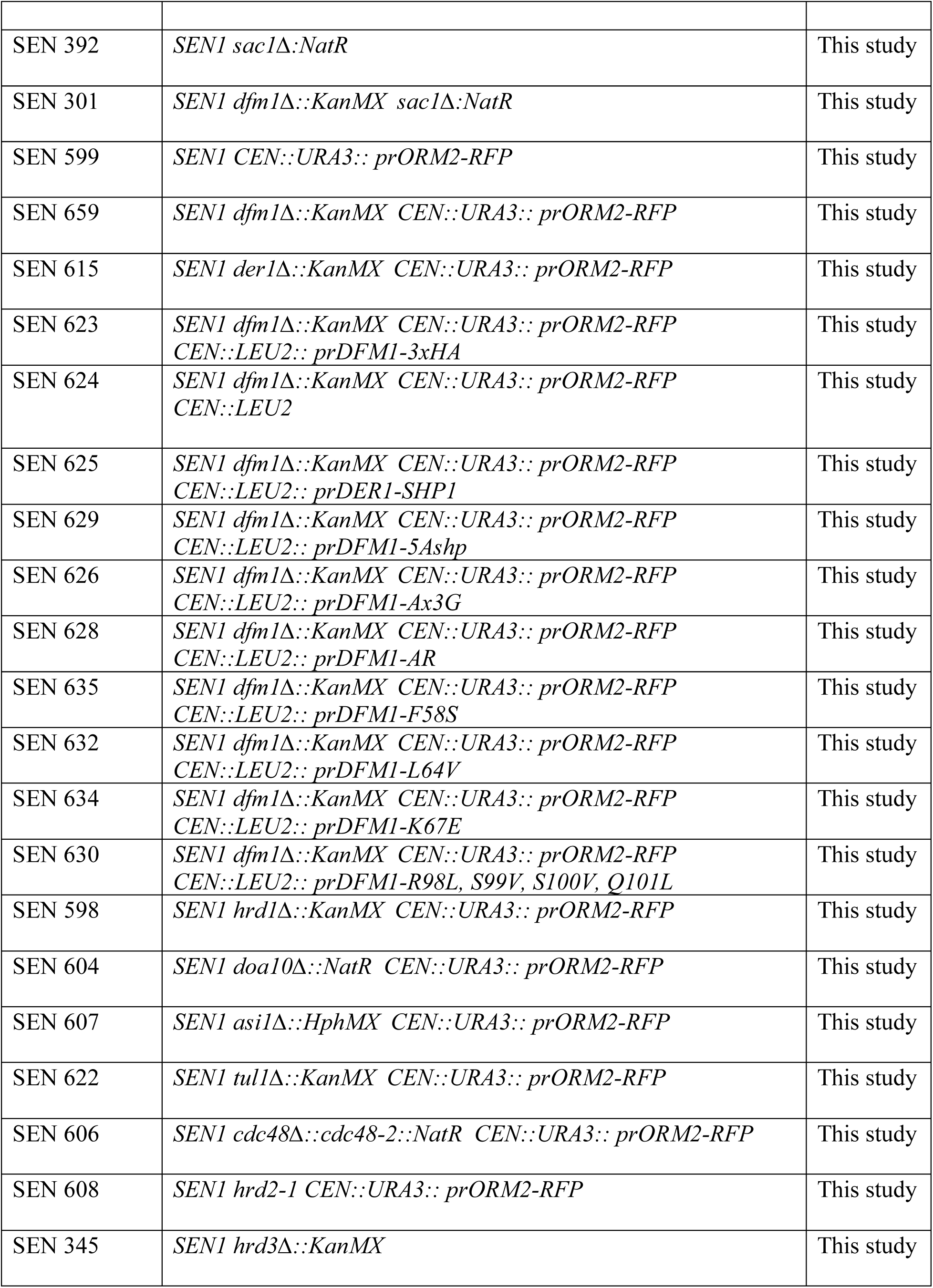
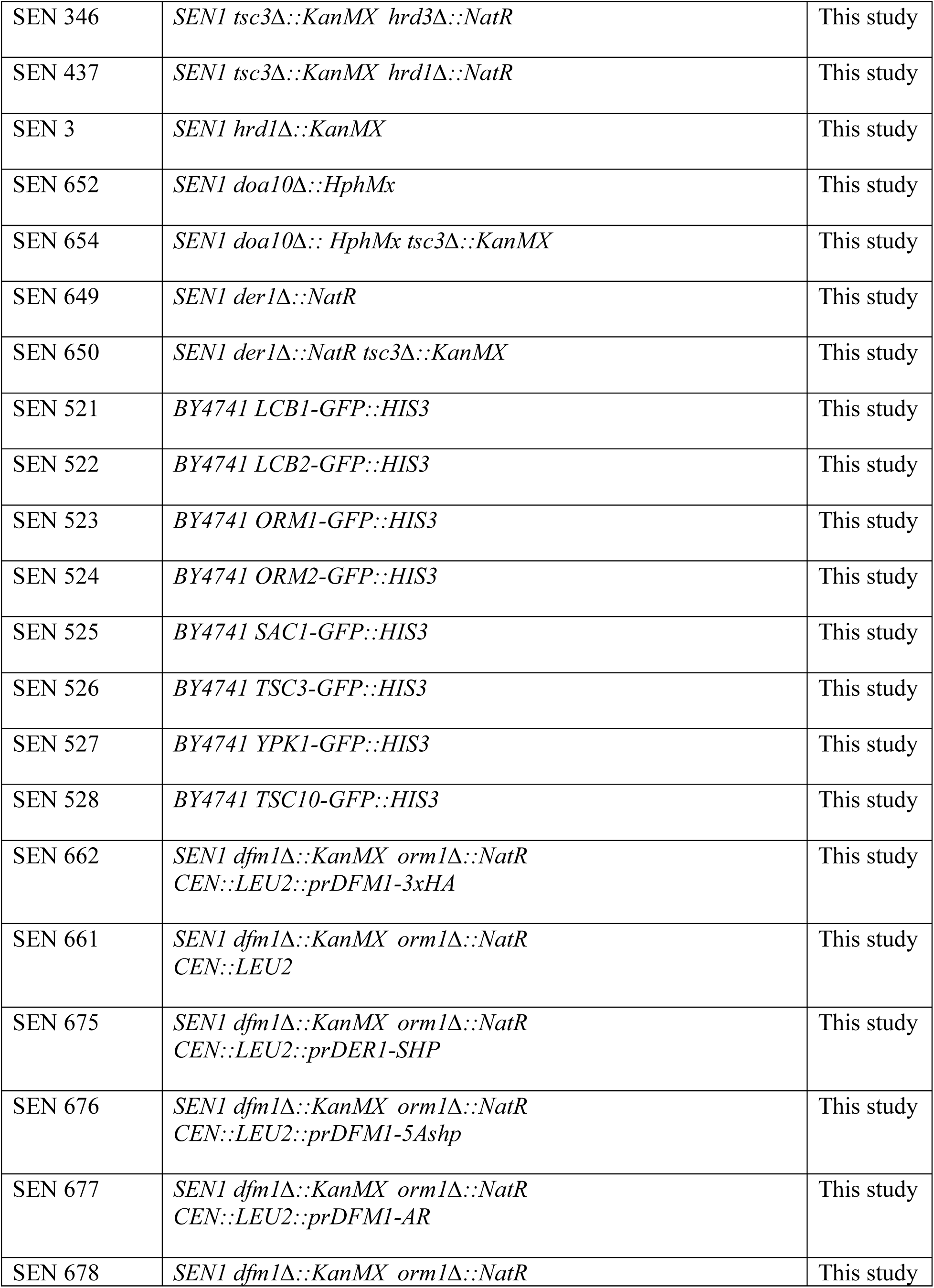
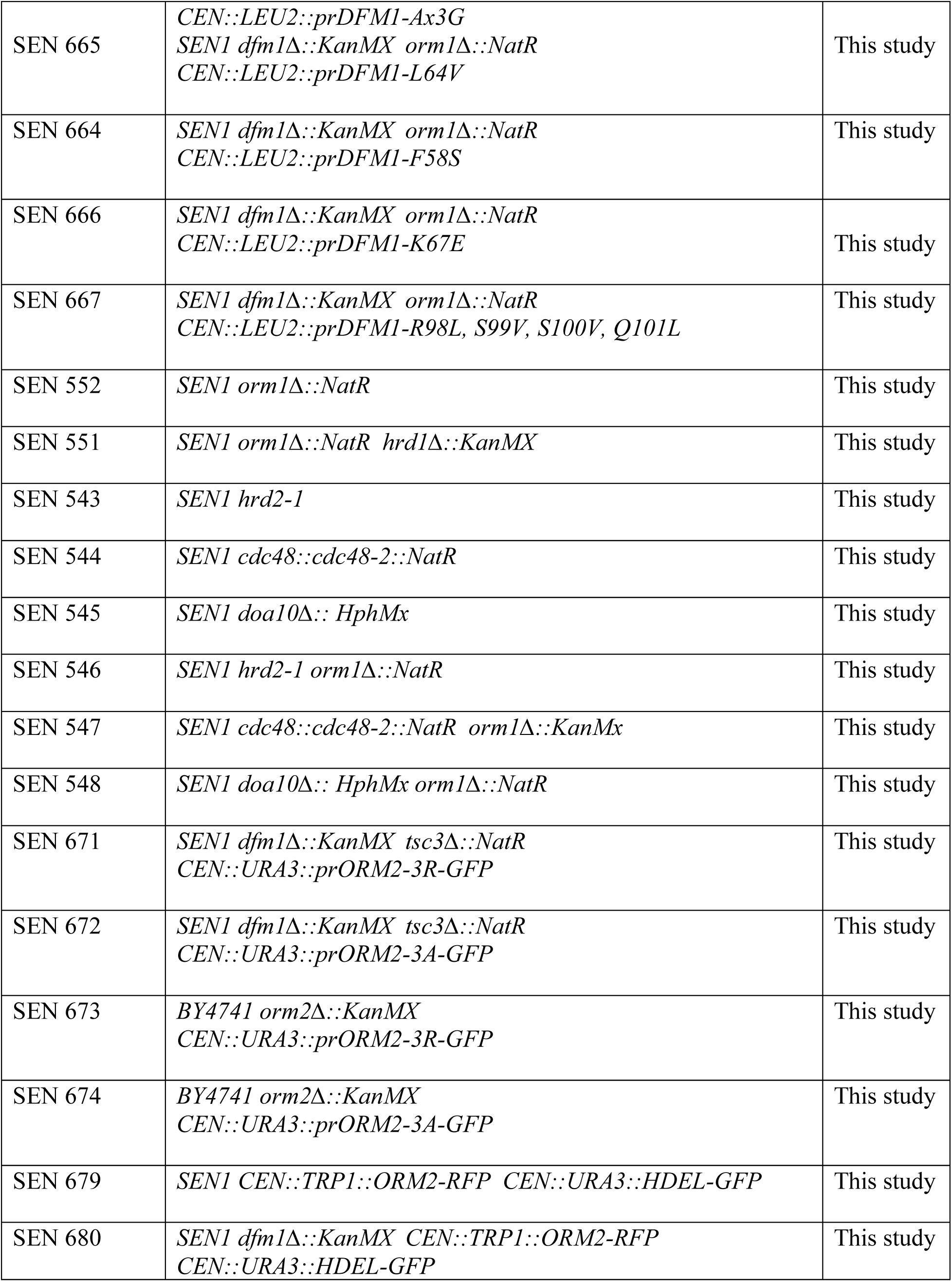
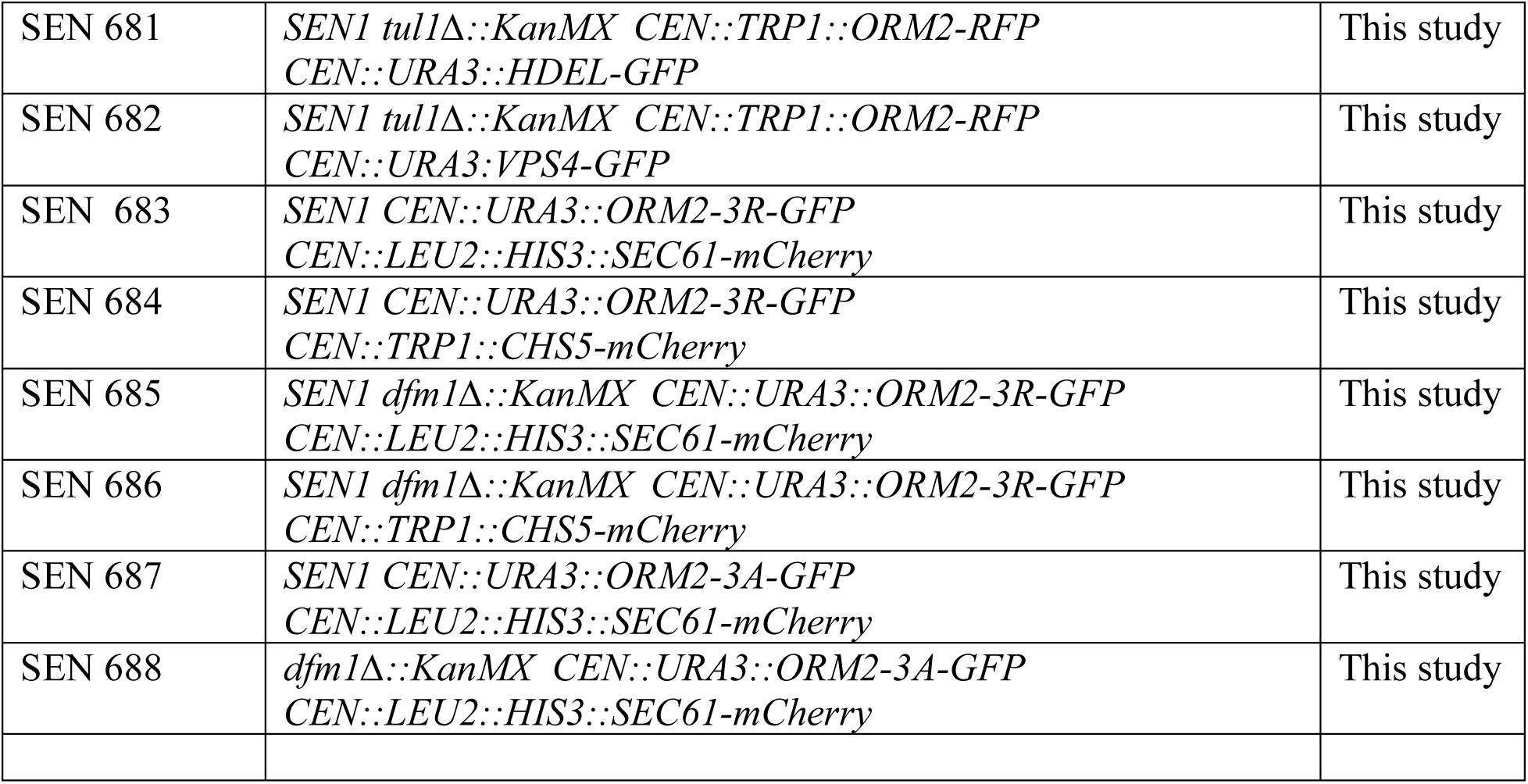
Yeast strains used in this study, Related to Figures 1–7

| **Strain** | **Genotype** | **Reference** |
| --- | --- | --- |
| SEN 1 | *Mata ade2-101 met2 lys-801 ura3-52 trp1::hisG leu2 his3Δ200* | This study |
| BY4741 | *Mata met15Δ0 his3Δ1 leu2Δ0 ura3Δ0* | This study |
| SEN 322 | *SEN1 CEN::URA3::GAL1pr-Dfm1-BirA-FLAG* | This study |
| SEN 325 | *SEN1 CEN::URA3::GAL1pr- BirA-FLAG* | This study |
| SEN 280 | *BY4741 DFM1::DFM1-GFP::HIS* | This study |
| SEN 439 | *BY4741 DFM1::DFM1-GFP::HIS*<br>*CEN::URA3:: prORM2-RFP* | This study |
| SEN 438 | *BY4741 DFM1::DFM1-GFP::HIS*<br>*CEN::LEU2:: prLCB1-RFP* | This study |
| SEN 635 | *SEN1 dfm1Δ::KanMX* | This study |
| SEN 637 | *SEN1 tsc3Δ::NatR* | This study |
| SEN 638 | *SEN1 dfm1Δ::KanMX tsc3Δ::NatR* | This study |
| SEN 640 | *SEN1 orm1Δ::KanMX* | This study |
| SEN 641 | *SEN1 orm1Δ::KanMX tsc3Δ::NatR* | This study |
| SEN 644 | *SEN1 orm2Δ::KanMX* | This study |
| RHY 646 | *SEN1 orm2Δ::KanMX tsc3Δ::NatR* | This study |
| SEN 285 | *SEN1 dfm1Δ::KanMX orm1Δ::NatR* | This study |
| SEN 291 | *SEN1 dfm1Δ::KanMX orm2Δ::NatR* | This study |
| SEN 280 | *BY4741 orm1Δ::KanMX* | This study |
| SEN 281 | *BY4741 orm2Δ::KanMX* | This study |
| SEN 369 | *BY4741 orm1Δ::KanMX orm2Δ::LEU2* | This study |
| SEN 282 | *BY4741 Lcb2-DaMP* | This study |
| SEN 266 | *BY4741 dfm1Δ::KanMX Lcb2-DaMP* | This study |
| SEN 392 | *SEN1 sac1Δ::NatR* | This study |
| SEN 301 | *SEN1 dfm1Δ::KanMX sac1Δ::NatR* | This study |
| SEN 599 | *SEN1 CEN::URA3:: prORM2-RFP* | This study |
| SEN 659 | *SEN1 dfm1Δ::KanMX CEN::URA3:: prORM2-RFP* | This study |
| SEN 615 | *SEN1 der1Δ::KanMX CEN::URA3:: prORM2-RFP* | This study |
| SEN 623 | *SEN1 dfm1Δ::KanMX CEN::URA3:: prORM2-RFP*<br>*CEN::LEU2:: prDFM1-3xHA* | This study |
| SEN 624 | *SEN1 dfm1Δ::KanMX CEN::URA3:: prORM2-RFP*<br>*CEN::LEU2* | This study |
| SEN 625 | *SEN1 dfm1Δ::KanMX CEN::URA3:: prORM2-RFP*<br>*CEN::LEU2:: prDER1-SHP1* | This study |
| SEN 629 | *SEN1 dfm1Δ::KanMX CEN::URA3:: prORM2-RFP*<br>*CEN::LEU2:: prDFM1-5Ashp* | This study |
| SEN 626 | *SEN1 dfm1Δ::KanMX CEN::URA3:: prORM2-RFP*<br>*CEN::LEU2:: prDFM1-Ax3G* | This study |
| SEN 628 | *SEN1 dfm1Δ::KanMX CEN::URA3:: prORM2-RFP*<br>*CEN::LEU2:: prDFM1-AR* | This study |
| SEN 635 | *SEN1 dfm1Δ::KanMX CEN::URA3:: prORM2-RFP*<br>*CEN::LEU2:: prDFM1-F58S* | This study |
| SEN 632 | *SEN1 dfm1Δ::KanMX CEN::URA3:: prORM2-RFP*<br>*CEN::LEU2:: prDFM1-L64V* | This study |
| SEN 634 | *SEN1 dfm1Δ::KanMX CEN::URA3:: prORM2-RFP*<br>*CEN::LEU2:: prDFM1-K67E* | This study |
| SEN 630 | *SEN1 dfm1Δ::KanMX CEN::URA3:: prORM2-RFP*<br>*CEN::LEU2:: prDFM1-R98L, S99V, S100V, Q101L* | This study |
| SEN 598 | *SEN1 hrd1Δ::KanMX CEN::URA3:: prORM2-RFP* | This study |
| SEN 604 | *SEN1 doa10Δ::NatR CEN::URA3:: prORM2-RFP* | This study |
| SEN 607 | *SEN1 asi1Δ::HphMX CEN::URA3:: prORM2-RFP* | This study |
| SEN 622 | *SEN1 tul1Δ::KanMX CEN::URA3:: prORM2-RFP* | This study |
| SEN 606 | *SEN1 cdc48Δ::cdc48-2::NatR CEN::URA3:: prORM2-RFP* | This study |
| SEN 608 | *SEN1 hrd2-1 CEN::URA3:: prORM2-RFP* | This study |
| SEN 345 | *SEN1 hrd3Δ::KanMX* | This study |
| SEN 346 | *SEN1 tsc3Δ::KanMX hrd3Δ::NatR* | This study |
| SEN 437 | *SEN1 tsc3Δ::KanMX hrd1Δ::NatR* | This study |
| SEN 3 | *SEN1 hrd1Δ::KanMX* | This study |
| SEN 652 | *SEN1 doa10Δ::HphMx* | This study |
| SEN 654 | *SEN1 doa10Δ:: HphMx tsc3Δ::KanMX* | This study |
| SEN 649 | *SEN1 der1Δ::NatR* | This study |
| SEN 650 | *SEN1 der1Δ::NatR tsc3Δ::KanMX* | This study |
| SEN 521 | *BY4741 LCB1-GFP::HIS3* | This study |
| SEN 522 | *BY4741 LCB2-GFP::HIS3* | This study |
| SEN 523 | *BY4741 ORM1-GFP::HIS3* | This study |
| SEN 524 | *BY4741 ORM2-GFP::HIS3* | This study |
| SEN 525 | *BY4741 SAC1-GFP::HIS3* | This study |
| SEN 526 | *BY4741 TSC3-GFP::HIS3* | This study |
| SEN 527 | *BY4741 YPK1-GFP::HIS3* | This study |
| SEN 528 | *BY4741 TSC10-GFP::HIS3* | This study |
| SEN 662 | *SEN1 dfm1Δ::KanMX orm1Δ::NatR*<br>*CEN::LEU2::prDFM1-3xHA* | This study |
| SEN 661 | *SEN1 dfm1Δ::KanMX orm1Δ::NatR*<br>*CEN::LEU2* | This study |
| SEN 675 | *SEN1 dfm1Δ::KanMX orm1Δ::NatR*<br>*CEN::LEU2::prDER1-SHP* | This study |
| SEN 676 | *SEN1 dfm1Δ::KanMX orm1Δ::NatR*<br>*CEN::LEU2::prDFM1-5Ashp* | This study |
| SEN 677 | *SEN1 dfm1Δ::KanMX orm1Δ::NatR*<br>*CEN::LEU2::prDFM1-AR* | This study |
| SEN 678 | *SEN1 dfm1Δ::KanMX orm1Δ::NatR* | This study |
| SEN 665 | *CEN::LEU2::prDFM1-Ax3G*<br>*SEN1 dfm1Δ::KanMX orm1Δ::NatR*<br>*CEN::LEU2::prDFM1-L64V* | This study |
| SEN 664 | *SEN1 dfm1Δ::KanMX orm1Δ::NatR*<br>*CEN::LEU2::prDFM1-F58S* | This study |
| SEN 666 | *SEN1 dfm1Δ::KanMX orm1Δ::NatR*<br>*CEN::LEU2::prDFM1-K67E* | This study |
| SEN 667 | *SEN1 dfm1Δ::KanMX orm1Δ::NatR*<br>*CEN::LEU2::prDFM1-R98L, S99V, S100V, Q101L* | This study |
| SEN 552 | *SEN1 orm1Δ::NatR* | This study |
| SEN 551 | *SEN1 orm1Δ::NatR hrd1Δ::KanMX* | This study |
| SEN 543 | *SEN1 hrd2-1* | This study |
| SEN 544 | *SEN1 cdc48::cdc48-2::NatR* | This study |
| SEN 545 | *SEN1 doa10Δ:: HphMx* | This study |
| SEN 546 | *SEN1 hrd2-1 orm1Δ::NatR* | This study |
| SEN 547 | *SEN1 cdc48::cdc48-2::NatR orm1Δ::KanMx* | This study |
| SEN 548 | *SEN1 doa10Δ:: HphMx orm1Δ::NatR* | This study |
| SEN 671 | *SEN1 dfm1Δ::KanMX tsc3Δ::NatR*<br>*CEN::URA3::prORM2-3R-GFP* | This study |
| SEN 672 | *SEN1 dfm1Δ::KanMX tsc3Δ::NatR*<br>*CEN::URA3::prORM2-3A-GFP* | This study |
| SEN 673 | *BY4741 orm2Δ::KanMX*<br>*CEN::URA3::prORM2-3R-GFP* | This study |
| SEN 674 | *BY4741 orm2Δ::KanMX*<br>*CEN::URA3::prORM2-3A-GFP* | This study |
| SEN 679 | *SEN1 CEN::TRP1::ORM2-RFP CEN::URA3::HDEL-GFP* | This study |
| SEN 680 | *SEN1 dfm1Δ::KanMX CEN::TRP1::ORM2-RFP*<br>*CEN::URA3::HDEL-GFP* | This study |
| SEN 681 | *SEN1 tul1Δ::KanMX CEN::TRP1::ORM2-RFP*<br>*CEN::URA3::HDEL-GFP* | This study |
| SEN 682 | *SEN1 tul1Δ::KanMX CEN::TRP1::ORM2-RFP*<br>*CEN::URA3::VPS4-GFP* | This study |
| SEN 683 | *SEN1 CEN::URA3::ORM2-3R-GFP*<br>*CEN::LEU2::HIS3::SEC61-mCherry* | This study |
| SEN 684 | *SEN1 CEN::URA3::ORM2-3R-GFP*<br>*CEN::TRP1::CHS5-mCherry* | This study |
| SEN 685 | *SEN1 dfm1Δ::KanMX CEN::URA3::ORM2-3R-GFP*<br>*CEN::LEU2::HIS3::SEC61-mCherry* | This study |
| SEN 686 | *SEN1 dfm1Δ::KanMX CEN::URA3::ORM2-3R-GFP*<br>*CEN::TRP1::CHS5-mCherry* | This study |
| SEN 687 | *SEN1 CEN::URA3::ORM2-3A-GFP*<br>*CEN::LEU2::HIS3::SEC61-mCherry* | This study |
| SEN 688 | *dfm1Δ::KanMX CEN::URA3::ORM2-3A-GFP*<br>*CEN::LEU2::HIS3::SEC61-mCherry* | This study |

**Table S3.**
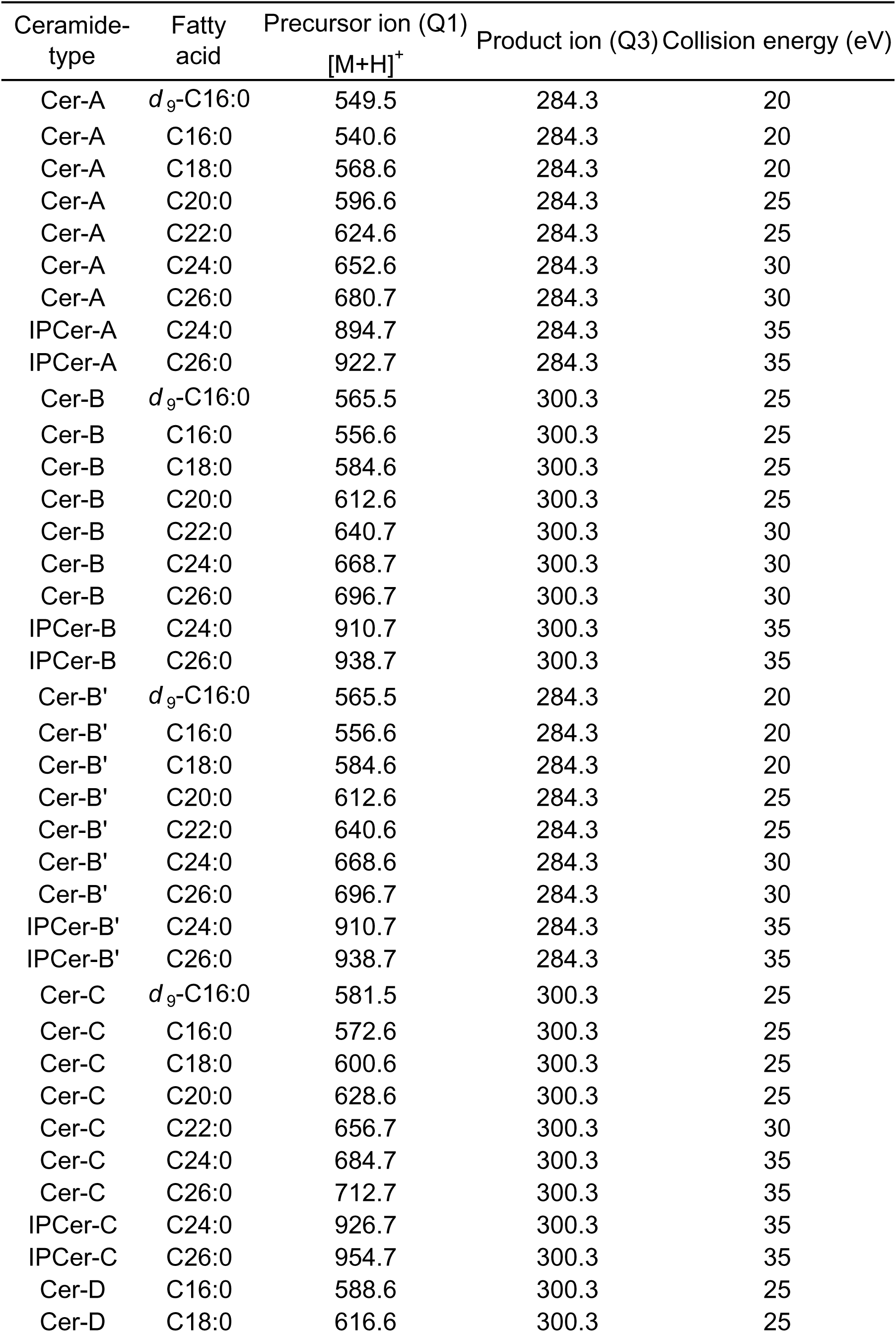
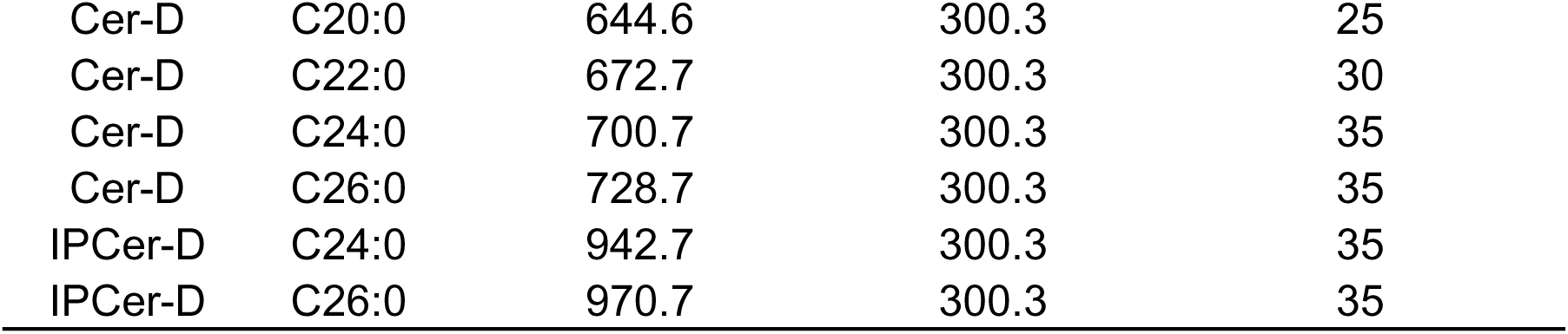
The values of m/z and collision energy for detection of ceramide species in LC-MS/MS analyses

| Ceramide-type | Fatty acid | Precursor ion (Q1)<br>[M+H]<sup>+</sup> | Product ion (Q3) | Collision energy (eV) |
| --- | --- | --- | --- | --- |
| Cer-A | *d*<sub>9</sub>-C16:0 | 549.5 | 284.3 | 20 |
| Cer-A | C16:0 | 540.6 | 284.3 | 20 |
| Cer-A | C18:0 | 568.6 | 284.3 | 20 |
| Cer-A | C20:0 | 596.6 | 284.3 | 25 |
| Cer-A | C22:0 | 624.6 | 284.3 | 25 |
| Cer-A | C24:0 | 652.6 | 284.3 | 30 |
| Cer-A | C26:0 | 680.7 | 284.3 | 30 |
| IPCer-A | C24:0 | 894.7 | 284.3 | 35 |
| IPCer-A | C26:0 | 922.7 | 284.3 | 35 |
| Cer-B | *d*<sub>9</sub>-C16:0 | 565.5 | 300.3 | 25 |
| Cer-B | C16:0 | 556.6 | 300.3 | 25 |
| Cer-B | C18:0 | 584.6 | 300.3 | 25 |
| Cer-B | C20:0 | 612.6 | 300.3 | 25 |
| Cer-B | C22:0 | 640.7 | 300.3 | 30 |
| Cer-B | C24:0 | 668.7 | 300.3 | 30 |
| Cer-B | C26:0 | 696.7 | 300.3 | 30 |
| IPCer-B | C24:0 | 910.7 | 300.3 | 35 |
| IPCer-B | C26:0 | 938.7 | 300.3 | 35 |
| Cer-B' | *d*<sub>9</sub>-C16:0 | 565.5 | 284.3 | 20 |
| Cer-B' | C16:0 | 556.6 | 284.3 | 20 |
| Cer-B' | C18:0 | 584.6 | 284.3 | 20 |
| Cer-B' | C20:0 | 612.6 | 284.3 | 25 |
| Cer-B' | C22:0 | 640.6 | 284.3 | 25 |
| Cer-B' | C24:0 | 668.6 | 284.3 | 30 |
| Cer-B' | C26:0 | 696.7 | 284.3 | 30 |
| IPCer-B' | C24:0 | 910.7 | 284.3 | 35 |
| IPCer-B' | C26:0 | 938.7 | 284.3 | 35 |
| Cer-C | *d*<sub>9</sub>-C16:0 | 581.5 | 300.3 | 25 |
| Cer-C | C16:0 | 572.6 | 300.3 | 25 |
| Cer-C | C18:0 | 600.6 | 300.3 | 25 |
| Cer-C | C20:0 | 628.6 | 300.3 | 25 |
| Cer-C | C22:0 | 656.7 | 300.3 | 30 |
| Cer-C | C24:0 | 684.7 | 300.3 | 35 |
| Cer-C | C26:0 | 712.7 | 300.3 | 35 |
| IPCer-C | C24:0 | 926.7 | 300.3 | 35 |
| IPCer-C | C26:0 | 954.7 | 300.3 | 35 |
| Cer-D | C16:0 | 588.6 | 300.3 | 25 |
| Cer-D | C18:0 | 616.6 | 300.3 | 25 |
| Cer-D | C20:0 | 644.6 | 300.3 | 25 |
| Cer-D | C22:0 | 672.7 | 300.3 | 30 |
| Cer-D | C24:0 | 700.7 | 300.3 | 35 |
| Cer-D | C26:0 | 728.7 | 300.3 | 35 |
| IPCer-D | C24:0 | 942.7 | 300.3 | 35 |
| IPCer-D | C26:0 | 970.7 | 300.3 | 35 |

